# Genome editing in plants using the compact editor CasΦ

**DOI:** 10.1101/2022.10.31.514567

**Authors:** Zheng Li, Zhenhui Zhong, Zhongshou Wu, Patrick Pausch, Basem Al-Shayeb, Jasmine Amerasekera, Jennifer A. Doudna, Steven E. Jacobsen

## Abstract

CRISPR-Cas systems have been developed as important tools for plant genome engineering. Here, we demonstrate that the hypercompact CasΦ nuclease is able to generate stably inherited gene edits in *Arabidopsis*, and that CasΦ guide RNAs can be expressed with either the Pol-III U6 promoter or a Pol-II promoter together with ribozyme mediated RNA processing. Using the *Arabidopsis fwa* epiallele we show that CasΦ displays higher editing efficiency when the target locus is not DNA methylated, suggesting that CasΦ is sensitive to chromatin environment. Importantly, two CasΦ protein variants, vCasΦ and nCasΦ, both showed much higher editing efficiency relative to the wildtype CasΦ enzyme, and yielded more offspring plants with inherited edits. Extensive genomic analysis of gene edited plants showed no off-target editing, suggesting that CasΦ is highly specific. The hypercompact size, T-rich minimal PAM and wide range of working temperatures make CasΦ an excellent supplement to existing plant genome editing systems.

**Significance Statement:** Plant genome engineering with CRISPR-Cas systems is frequently used in both research and agriculture. Here, we demonstrate that the hypercompact CasΦ-2 nuclease is able to generate heritable gene edits in *Arabidopsis*. Two CasΦ protein variants vCasΦ and nCasΦ increased the editing efficiency in plants. CasΦ also has a wide range of working temperatures and the editing by CasΦ is highly specific. We also observed that editing by CasΦ is sensitive to chromatin environment. The hypercompact size, T-rich minimal PAM and wide range of working temperatures make CasΦ an excellent supplement to existing plant genome editing systems.

## Introduction

CRISPR-Cas systems, originally discovered as an adaptive immune system in bacteria and archaea (1–3), have been widely used as genome engineering tools (4–6). Class 2 CRISPR-Cas systems are of particular interest as the guide RNA binding and the cleavage of target nucleic acids are accomplished by a single effector protein (7). For plant genome engineering, Cas9 (2, 8–12) and Cas12a (13–17) have been routinely applied in multiple plant species. With the discovery of novel CRISPR-Cas systems, additional DNA and RNA targeting Cas proteins serve as potential plant genome engineering tools. CRISPR-CasΦ, discovered from huge bacteriophages (18), targets double stranded DNA and generates staggered cuts (19). Importantly, CasΦ proteins are only 700 to 800 amino acids (aa) (19), which are much smaller in size compared to Cas9 (1000-1400 aa) (2, 20, 21) and Cas12a (1100-1300 aa) (13). The compact sizes of the CRISPR-CasΦ systems may allow for approaches where protein or nucleic acid size is a limiting factor. CasΦ systems also have T rich minimal PAM sequences (19). Huge bacteriophages encoding CasΦ systems are from diverse ecosystems potentially providing a range of optimum temperatures for CasΦ activity (18). CasΦ proteins are therefore interesting candidates for novel plant genome engineering tools.

The CasΦ-2 ortholog was previously shown to be capable of target DNA editing in both human and plant cells (19). It recognizes a 5’-TBN-3’ PAM sequence (where B is G, T or C) and generates staggered 5’ overhangs which usually yield multiple base pair deletions after the action of cellular DNA repair machineries (19). Similar to Cas12a, the CasΦ-2 protein is also able to process pre-crRNA into mature crRNAs (19). The CasΦ-2 protein employs a single RuvC active site for DNA cleavage, which is also used for pre-crRNA processing (19, 22). A structural study of CasΦ-2 revealed that helix α7 blocks the path of the target strand of the substrate DNA towards a PAM-proximal position (22). Mutations in the negatively charged tip of helix α7 either by substitution of the whole negatively charged tip to a linker sequence GSSG (vCasΦ) or substitution of residues in the negatively charged tip to alanine (E159A, S160A, S164A, D167A, E168A) (nCasΦ) resulted in significantly faster substrate DNA cleavage compared to the wild-type (WT) CasΦ (22). This finding provides potentially more efficient CasΦ variants for genome engineering purposes.

In this study, we provide evidence that CasΦ can be utilized as a novel plant genome editing tool to generate heritable mutations. The CasΦ genome editing system is compatible with Pol-II promoter driven guide RNA transcription and ribozyme mediated guide RNA processing machineries. We also found that CasΦ editing is more efficient at unmethylated DNA than at methylated DNA. Finally, the hyperactive CasΦ variants vCasΦ and nCasΦ exhibited much higher editing efficiency with all of the gRNAs tested, and off-target editing of the CasΦ variants vCasΦ and nCasΦ was not observed, suggesting that these variants can be utilized for highly specific genome editing in plants.

## Results

### CasΦ-2 is capable of generating heritable mutations in *Arabidopsis*

Previously, it was shown that CasΦ-2 RNPs were capable of editing the *AtPDS3* gene in *Arabidopsis* mesophyll protoplasts (19). To investigate if CasΦ-2 is able to edit a target gene in transgenic plants, we used the *Arabidopsis UBQ10* promoter to drive the expression of an *Arabidopsis* codon optimized CasΦ-2 and the U6 promoter to drive transcription of guide RNAs (Fig. 1*A*). The ubiquitous expression by the *UBQ10* promoter allows for assessment of editing efficiency in somatic tissues of transgenic plants as well as using mesophyll protoplasts for further optimization. We utilized *AtPDS3* gRNA10, which had the highest editing efficiency of all *AtPDS3* gRNAs tested in protoplasts (19). Version 1 and version 2 constructs were made with nuclear localization signal peptides either flanking N- and C-termini or only at the C-terminus of the CasΦ-2 protein, respectively (Fig. 1*A*). Seventy-eight T1 transgenic plants of the version 1 construct were screened by Sanger sequencing and a plant heterozygous for mutation of the gRNA10 targeted region was identified (T1 #33) (Fig. 1*B*). Further amplicon sequencing of different parts of this plant indicated that it was mosaic for the heterozygous mutation, with leaf 1 and leaf 2 showing about 50% of reads carrying mutation, but other leaves showing little to no editing (Fig. 1*C*). The dominant form of mutation detected in this plant by amplicon sequencing was a 6 bp deletion in the *AtPDS3* gR10 targeted region, although a small number of reads with other forms of deletion were also detected (Table S1). The *AtPDS3* gene encodes a phytoene desaturase enzyme that is essential for chloroplast development. Disruption of this gene function results in albino and dwarfed seedlings (23). Indeed, albino and dwarfed seedlings were observed from the offspring (T2) population of plant #33 (Fig. 1*D*). 20 albino/dwarf seedlings were tested individually for the *AtPDS3* gR10 target region and all of the tested seedlings were homozygous for a 6 bp deletion at the gR10 target region (Fig. 1*E*). This 6 bp deletion leads to the loss of two amino acids which are highly conserved among orthologous proteins (Fig. S1*A*). In addition, two of these 20 albino/dwarf T2 seedlings had segregated away the transgene and were thus CasΦ-2 transgene free (Fig. 1*F*), confirming the germline transmission (heritability) of the CasΦ-2 generated mutation in the *AtPDS3* gene from the T1 to the T2 plants.

**Fig. 1.**
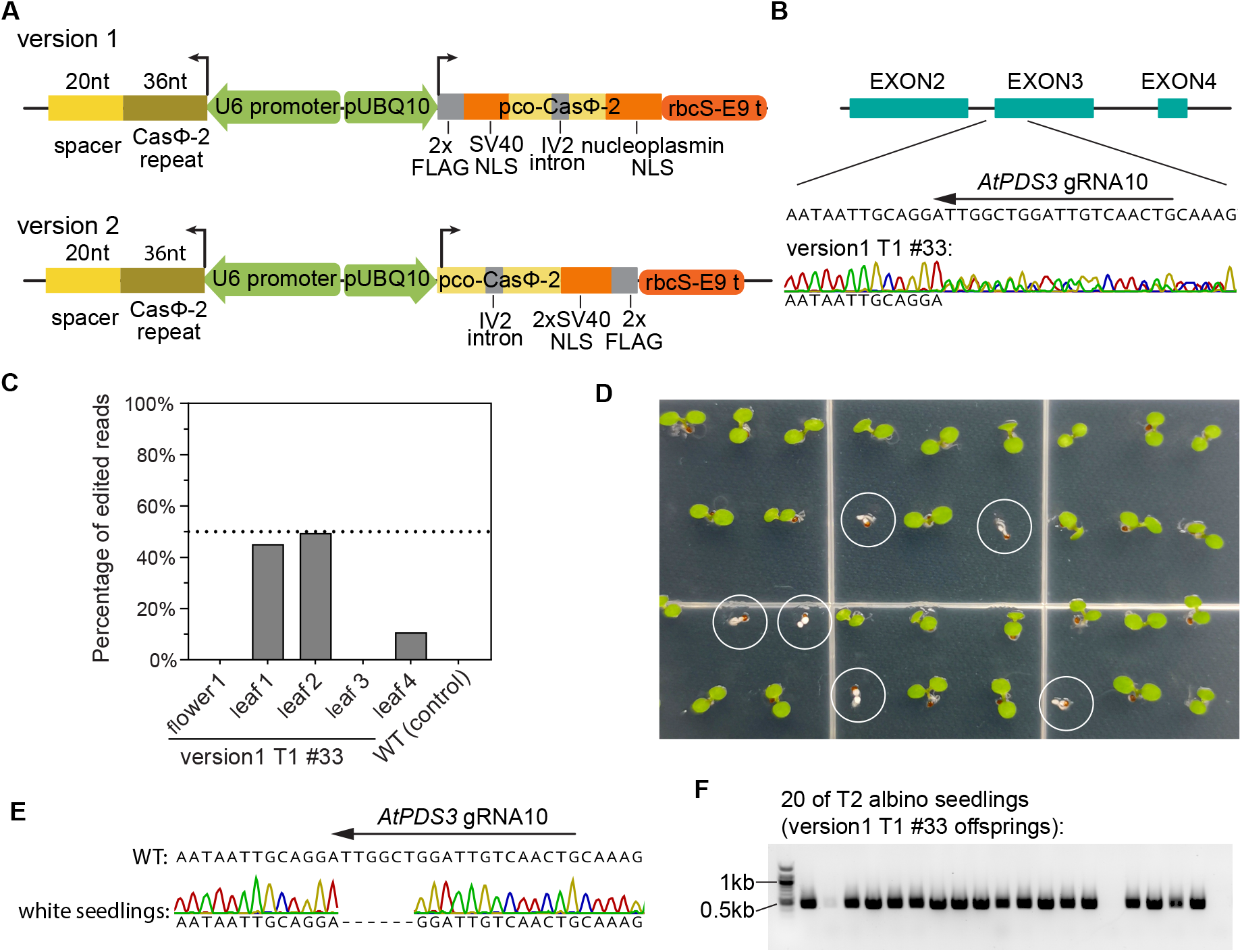
Editing of the *AtPDS3* gene by CasΦ-2 in transgenic plants. **(A)** Schematics of the version 1 and version 2 constructs to generate *Arabidopsis* transgenic plants expressing the CasΦ-2 protein and the CasΦ-2 guide RNA. NLS, nuclear localization signal. pUBQ10, *Arabidopsis UBQ10* gene promoter. pco-CasΦ-2, *Arabidopsis* codon optimized CasΦ-2. rbcS-E9 t, rbcS E9 terminator. U6 promoter, *AtU6-26* gene promoter. **(B)** Sanger sequencing result of T1 transgenic plant #33 of version 1 construct with *AtPDS3* gRNA10 at the target region. **(C)** Amplicon sequencing results of different parts of T1 plant #33 containing version 1 construct for the *AtPDS3* gRNA10 targeted region. **(D)** Picture of seedlings from T2 populations of T1 plant #33, with albino seedlings circled. **(E)** Sanger sequencing result of albino seedlings from the T2 populations of T1 plant #33 at the *AtPDS3* gRNA10 targeted region. (**F)** PCR amplification of the DNA of 20 randomly selected albino seedlings from the T2 population of T1 plant #33 for a fragment of the CasΦ-2 transgene.

For the version 2 construct, six T1 transgenic plants were screened, this time directly by amplicon sequencing so that we could more sensitively detect plants with low editing efficiency. T1 plant #6 showed the highest number of edits, with 13.5% of reads carrying mutations, while the other five plants showed much lower or no editing (Fig. S1*B*). Ninety-six offspring T2 plants of T1 plant #6 were analyzed and six of them were heterozygous for mutations at the *AtPDS3* gR10 targeted region (Fig. S1*C*). As expected, albino seedlings were identified in the T3 offspring populations of these heterozygous T2 plants (Fig. S1*D*). DNA sequencing of T3 albino seedlings derived from different T2 lineages showed deletions of 3 bp, 5 bp, 9 bp (two plants), or a 42 bp deletion plus a 1 bp deletion that was 16 bp upstream of the larger deletion (two plants) (Fig. S2*A*). CasΦ-2 transgene free albino seedlings were also identified from these T3 populations, further supporting the heritability of the CasΦ-2 generated mutation in the *AtPDS3* gene (Fig. S2*B*). These results suggest that the initial T1 plant #6 was chimeric for different deletions which were inherited into the different T2 and T3 lines.

### RDR6 mediated transgene silencing attenuates CasΦ-2 mediated editing in transgenic plants

In *Arabidopsis*, RNA-dependent RNA polymerase 6 (RDR6) plays an important role in the initiation of transgene silencing (24, 25). To evaluate if the CasΦ-2 transgene is also affected by transgene silencing, the editing efficiencies of CasΦ-2 transgenic T1 plants in the WT background or in the *rdr6-15* mutant (26) background were compared. For both the version 1 and the version 2 constructs with *AtPDS3* gRNA10, we detected significantly higher editing efficiency in the population of T1 transgenic plants in the *rdr6-15* mutant background compared to the WT background (Fig. S3*A*), suggesting that RDR6 mediated silencing is limiting the editing efficiency in CasΦ-2 transgenic plants. No significant difference was detected between the version 1 and version 2 constructs in the same genetic background, suggesting that the two configurations of nuclear localization signal work similarly (Fig. S3*A*).

### CasΦ-2 shows similar editing efficiency at 23 °C and 28 °C

We previously observed that CasΦ-2 RNPs with gRNA10 produced around 0.85% edits in *Arabidopsis* protoplasts using a 2-day room temperature incubation with a 2.5-hour 37 °C pulse (79,768 edited reads/9,404,589 total reads) (19). We performed a similar experiment in the absence of the 37 °C pulse, and found similar (1.1%) efficiency (91,950 edited reads/8,572,470 total reads), suggesting that CasΦ-2 is functional at lower temperatures. *Arabidopsis* grows ideally at around 23 °C but can also grow at temperatures up to about 28 °C (27), so we also performed an experiment with the version 2 plasmid comparing editing efficiencies in protoplasts at 23 °C vs 28 °C and found a roughly similar editing efficiency in both with gRNA10 (roughly 1.1%) (Fig. S3*B*). To test the effect of temperature with a longer incubation time and in plants stably expressing CasΦ-2, T1 transgenic plants of version 1 or version 2 constructs either in the wild type Col-0 background or the *rdr6-15* background were either continuously grown at room temperature (23 °C) or initially grown at 28 °C for two weeks then transferred to room temperature. No significant difference was detected between the editing efficiencies of the T1 plants grown under these two temperature regimes (Fig. S3 *C and D*), suggesting that CasΦ-2 expressing transgenes function similarly between 23 °C and 28 °C. Previously, CasΦ-2 has also been shown to be functional at 37 °C in human cells (19). Altogether, these data suggests that the CasΦ-2 protein has a wide functional temperature range, which should make it useful for editing in a wide variety of eukaryotic systems.

### CasΦ-2 mediated editing is sensitive to DNA methylation status

To study the effect of chromatin compaction status on the efficiency of CasΦ-2 editing, we took advantage of the *FWA* gene and the *fwa-4* epiallele. In wild type (WT) *Arabidopsis* plants, the *FWA* gene is silent in all adult plant tissues owing to DNA methylation in the promoter. *FWA* is normally only expressed by the maternal allele in the developing endosperm where it is imprinted and demethylated (28). In the epiallele *fwa-4*, the promoter is heritably unmethylated and thus the *FWA* gene is expressed ectopically (29). Ten guide RNAs were designed targeting the promoter region of the *FWA* gene, immediately upstream of the transcription start site (Fig. 2*A*). In WT plants, this region contains DNA methylation and is targeted by the RNA directed DNA methylation machinery (30, 31), as evidenced by Pol V occupancy (Fig. 2*A*). ATAC-seq signals are also significantly lower in the WT plant compared to the *fwa-4* epi-mutant (32) (Fig. 2*A*), suggesting that the accessibility of the *FWA* promoter region is lower in the WT plant. An *in vitro* cleavage assay with CasΦ-2 RNPs showed that all 10 *FWA* guide RNAs led to cleavage of the PCR amplified *FWA* gene fragment, with the majority of the substrate DNA cleaved by RNPs of gRNA1, gRNA4, gRNA5, gRNA6, and gRNA7 (Fig. S4). When CasΦ-2 RNPs were transfected into *fwa-4* epi-mutant protoplasts, gene editing events were detected with gRNA1, gRNA4, gRNA5 and gRNA6 (Fig. 2*B*). To compare the editing efficiency at *FWA* in different chromatin states, in an independent experiment, protoplasts of WT and *fwa-4* epi-mutant plants were prepared and transfected with CasΦ-2 RNPs with *FWA* gRNA1, gRNA4, gRNA5 and gRNA6. Much higher editing efficiency was observed for each of the four gRNAs in the *fwa-4* protoplasts compared to the WT protoplasts (Fig. 2*C*), suggesting that the CasΦ-2 mediated editing is more efficient when the target DNA has a more open chromatin state compared to a repressive chromatin state.

**Fig. 2.**
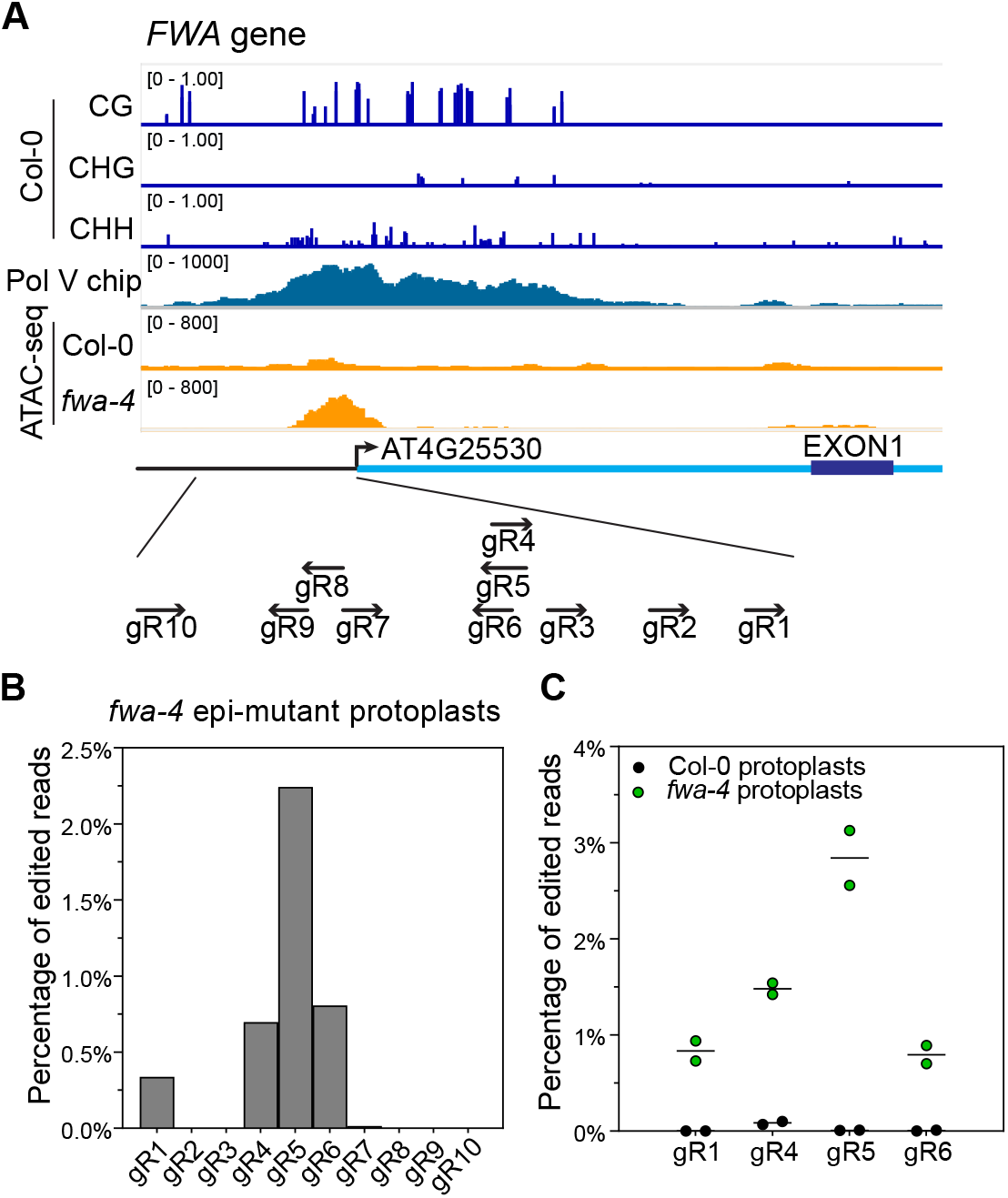
Effect of the DNA methylation status at the *FWA* gene promoter on editing efficiency by CasΦ-2. (A) DNA methylation level and normalized Pol V chip signal (normalized by RPKM) in Col-0 plants, as well as normalized ATAC-seq signal (normalized by RPKM) in Col-0 plants versus the *fwa-4* epi-mutant plants at the *FWA* gene (AT4G25530) promoter region. The relative positions of the CasΦ-2 guide RNAs are illustrated in the schematic below. (B) CasΦ-2 editing efficiencies at the target loci in the *FWA* promoter region with gRNA 1 to gRNA 10 in the *fwa-4* epi-mutant protoplasts. (C) Comparison of CasΦ-2 editing efficiencies in the *FWA* promoter region with *FWA* gRNA1, gRNA4, gRNA5 and gRNA6 in Col-0 protoplasts versus in *fwa-4* epi-mutant protoplasts. Two replicates were performed for each gRNA in each type of protoplast. Dots indicate the editing efficiencies of individual replicates and the line indicates the mean of the two replicates.

### CasΦ-2 guide RNAs can be expressed with Pol-II promoters and ribozyme processing machineries

As Pol-II promoters have been successfully used to drive the gRNA transcription for the Cas9 and Cas12 systems (14, 33), we wanted to test if Pol II promoters are able to drive CasΦ-2 guide RNA transcription. Three Pol II promoter and terminator sets were tested, the CmYLCV promoter and 35S terminator, a 2×35S promoter and HSP18.2 terminator, and a UBQ10 promoter with a rbcS-E9 terminator (Fig. 3*A*). Since CasΦ-2 has intrinsic pre-crRNA processing activity (19), *AtPDS3* gRNA10 without additional RNA processing machinery in three configurations were tested, a single CasΦ-2 repeat followed by *AtPDS3* gRNA10 spacer, a CasΦ-2 repeat followed by *AtPDS3* gRNA10 spacer followed by a second CasΦ-2 repeat, and a triple array of CasΦ-2 repeats/*AtPDS3* gRNA10 spacers followed by a fourth CasΦ-2 repeat (Fig. 3*A*). Among the three combinations of Pol-II promoters and terminators, the CmYLCV promoter with the 35S terminator led to the highest editing efficiency, while the UBQ10 promoter with the rbCS-E9 terminator led to the lowest editing efficiency in protoplasts (Fig. 3*B*). Out of the three different gRNA configurations, the single CasΦ-2 repeat followed by the *AtPDS3* gRNA10 exhibited the highest editing efficiency, while the CasΦ-2 repeat followed by the *AtPDS3* gRNA10 with another CasΦ-2 repeat at the end exhibited the lowest editing efficiency (Fig. 3*B*). When combining the CmYLCV promoter/ 35S terminator with the single CasΦ-2 repeat followed by the *AtPDS3* gRNA10, the target gene editing efficiency was much higher than that of the AtU6-26 *AtPDS3* gRNA10 cassette in protoplasts (Fig. 3*B*). Consistent with the higher levels of editing observed, a higher level of *AtPDS3* gRNA10 was detected in protoplasts transfected with plasmid carrying the cassette with the CmYLCV promoter *AtPDS3* gRNA10 construct (Fig. S5). This data suggests that boosting the levels of gRNAs using a strong promoter can increase the efficiency of gene editing by CasΦ-2.

**Fig. 3.**
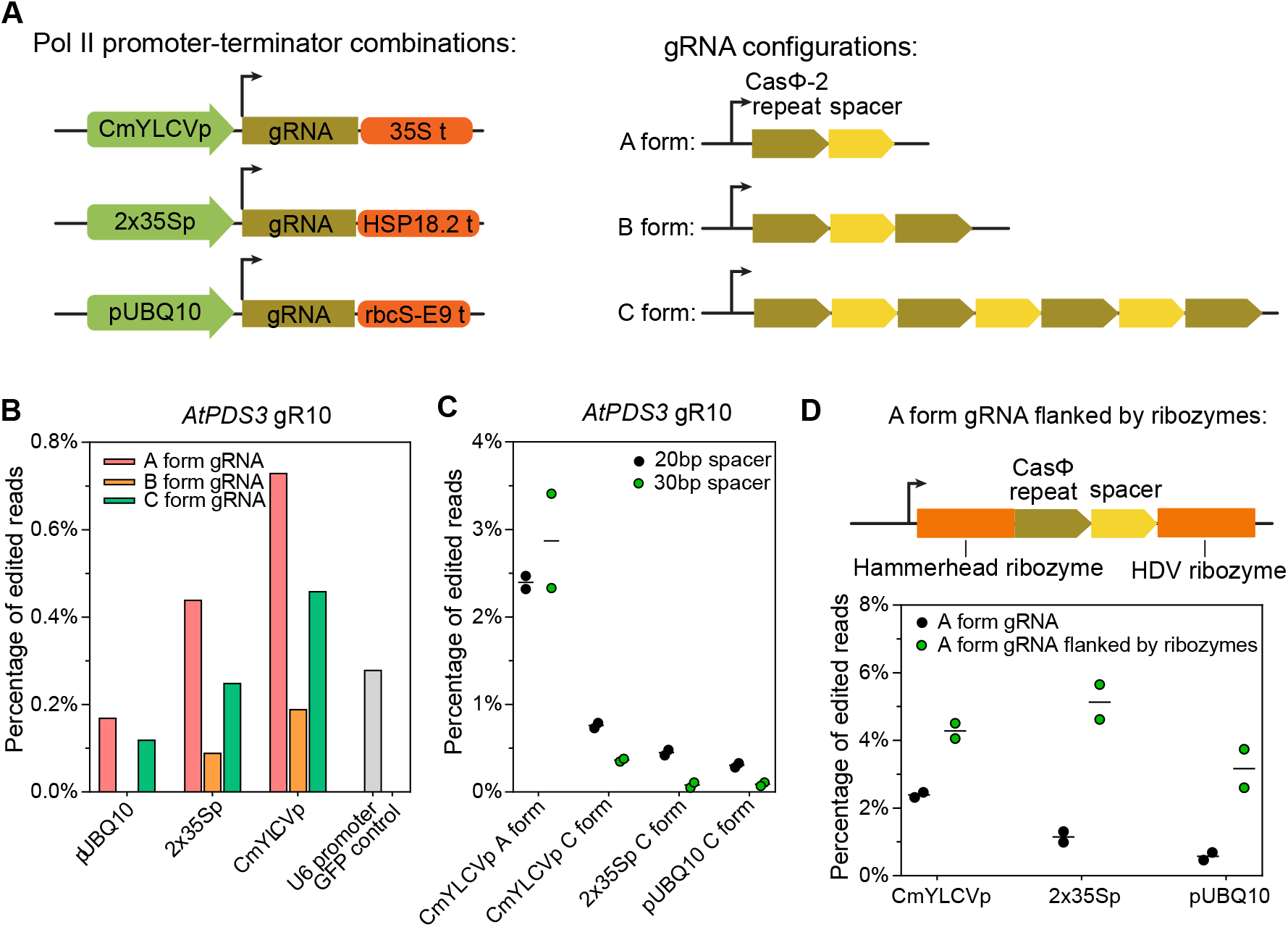
Pol-II promoter-terminator combinations and ribozymes for transcription and processing of CasΦ-2 guide RNA. (A) Schematics of the tested Pol-II promoter-terminator combinations (left panel) and the configurations of Cas Φ-2 guide RNA arrays (right panel). CmYLCVp, Cestrum yellow leaf curling virus promoter. 35S t, 35S terminator. 2×35Sp, two times 35S promoter. HSP18.2 t, *Arabidopsis HSP18.2* gene terminator. For gRNA configurations: A form, a single CasΦ-2 repeat followed by *AtPDS3* gRNA10 spacer; B form, a CasΦ-2 repeat followed by *AtPDS3* gRNA10 spacer followed by a second CasΦ-2 repeat; C form, a triple array of CasΦ-2 repeats/*AtPDS3* gRNA10 spacers followed by a fourth CasΦ-2 repeat. (B) Summary of the target (*AtPDS3*) editing efficiencies in protoplasts, comparing promoter-terminator combinations and gRNA configurations, with the U6 promoter driving *AtPDS3* gRNA10 as a control. (C) CasΦ-2 editing efficiencies of *AtPDS3* gRNA10 with 20 bp spacer and 30 bp spacer were compared. (D) CasΦ-2 editing efficiencies of *AtPDS3* gRNA10 with or without the ribozyme processing machineries were compared. For (C) and (D), individual replicate values and mean of the two replicates of each test were plotted.

The fact that the single *AtPDS3* gRNA10 without another CasΦ-2 repeat at the end exhibited the highest editing efficiency among the three gRNA configurations suggests that the CasΦ-2 processing of the transcript was not able to provide enough mature gRNAs. The 20 bp spacer could be too short for CasΦ-2 to bind simultaneously to both of the repeats flanking the spacer for pre-crRNA processing. To examine this further, *AtPDS3* gRNA10 with a 30 bp spacer was used to test if a longer spacer might assist the self-processing of pre-crRNA by CasΦ-2. However, we observed no difference between the editing efficiencies of the 30 bp and the 20 bp *AtPDS3* gRNA10 spacers when constructed in the A form (Fig. 3*C*). We also observe a slight decrease of target gene editing when the 30 bp spacer was constructed in the C form compared to the 20 bp spacer (Fig. 3*C*). These results suggest that a longer 30 bp spacer did not promote more efficient processing of pre-crRNA by CasΦ-2.

We tested whether adding a secondary gRNA processing mechanism might facilitate the release of mature gRNA. A ribozyme gRNA processing system, previously proven to be able to facilitate Cas9 and Cas12a guide RNA processing (14, 34), was cloned to flank the CasΦ-2 repeat and the *AtPDS3* gRNA10 spacer sequence. The Hammerhead (HH) type ribozyme was added to the 5’ end of the CasΦ-2 *AtPDS3* gRNA10 and a hepatitis delta virus (HD) ribozyme was added to the 3’ end (Fig. 3*D*). Constructs with ribozymes led to significantly higher editing efficiency, with all three promoter-terminator combinations tested (Fig. 3*D*). These results suggest that ribozymes were able to promote the processing of gRNA from the Pol II transcripts, leading to a higher editing efficiency.

### The engineered CasΦ-2 variants vCasΦ and nCasΦ have higher target gene editing efficiency in protoplasts

It has been previously reported that the engineered CasΦ-2 variants vCasΦ and nCasΦ cleaved substrate DNA faster than the WTCasΦ *in vitro* (22). To test if these engineered CasΦ-2 variants are able to edit a target gene with higher efficiency in *Arabidopsis*, mesophyll protoplasts were transfected with plasmids expressing WTCasΦ, vCasΦ, or nCasΦ, as well as the desired guide RNAs. Version 2 constructs with U6 promoter driven *AtPDS3* gRNA8 (19) and gRNA10 were transfected into WT protoplasts, while Version 2 constructs with U6 promoter driven *FWA* gRNA1, gRNA4, gRNA5 and gRNA6 were transfected into *fwa-4* protoplasts. Higher editing efficiencies were observed with vCasΦ and nCasΦ variants for all guide RNAs tested (Fig. 4*A*). To statistically evaluate the differences between the editing efficiencies, for the 6 gRNAs tested, normalized editing efficiencies were calculated (ratio over WTCasΦ efficiency) and pooled for statistical tests. Both the vCasΦ and nCasΦ variants yielded significantly higher editing efficiencies than the WTCasΦ, with ~17 fold of increase in editing efficiency for vCasΦ and ~10 fold of increase in editing efficiency for nCasΦ (Fig. 4*B*). Similar tests were performed with RNPs reconstituted with WTCasΦ, vCasΦ, or nCasΦ and the same guide RNAs (Fig. S6*A*). A significant increase in the editing efficiency was detected with both the vCasΦ and nCasΦ variants when pooled analysis was performed (Fig. S6*B*), although the overall fold increase in editing efficiency was lower compared to the plasmid transfection experiments (Fig. 4*B* and Fig. S6*B*).

**Fig. 4.**
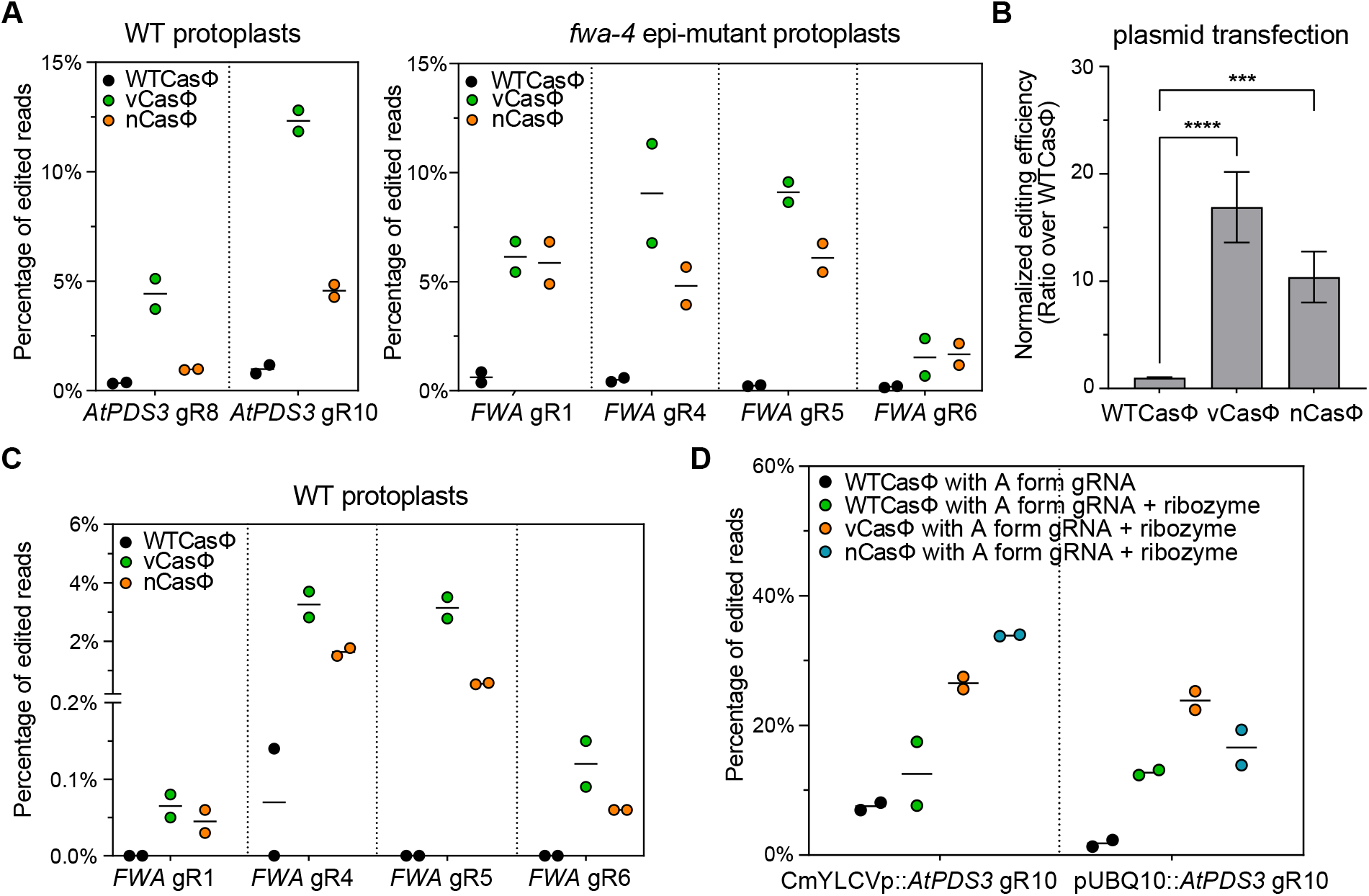
Comparison of editing efficiency by the vCasΦ and nCasΦ variants and WTCasΦ in protoplasts. (A) Plasmids with WTCasΦ, vCasΦ and nCasΦ expression cassettes and with indicated guide RNAs were transfected into protoplasts prepared from Col-0 plants (WT) (left panel) and from *fwa-4* epi-mutant plants (right panel). (B) Target gene editing efficiencies in (A) were normalized by calculating the ratio of editing efficiencies over that of mean editing efficiency by WTCasΦ for each guide RNA. Mean and standard error of the normalized editing efficiencies for all gRNAs were plotted. Unpaired t-test was used to calculate P value of indicated comparisons. ***, 0.0001<P<0.001, ****, P<0.0001. (C) Plasmids with WTCasΦ, vCasΦ and nCasΦ expression cassettes and guide RNAs targeting the *FWA* promoter region were transfected into WT protoplasts. (D) Plasmids with WTCasΦ, vCasΦ and nCasΦ expression cassettes and *AtPDS3* gRNA10 with or without ribozymes driven by CmYLCVp and pUBQ10 were transfected into protoplasts. For (A), (C) and (D), individual replicate values and mean of the two replicates of each test were plotted.

We also tested if the vCasΦ and nCasΦ variants were able to enhance the editing efficiency at more compact chromatin utilizing the *FWA* promoter region as the target. To test this, protoplasts from WT plants were used for plasmid transfections with *FWA* gRNA1, gRNA4, gRNA5 and gRNA6. Although zero to very low editing was observed for WTCasΦ, both vCasΦ and nCasΦ variants yielded readily detectable and higher editing frequencies for all four *FWA* guide RNAs tested (Fig. 4*C*). Thus, in compact chromatin environments, the vCasΦ and nCasΦ variants appear to dramatically enhance the editing efficiency compared to WTCasΦ.

To confirm that the vCasΦ and nCasΦ variants, like WTCasΦ, are compatible with guide RNAs driven by Pol-II promoters, the CmYLCV promoter and the UBQ10 promoter were used to drive transcription of *AtPDS3* gRNA10 flanked by ribozymes. For both the CmYLCV promoter and the UBQ10 promoter, the editing efficiencies of the vCasΦ and nCasΦ variants were higher than that of the WTCasΦ (Fig. 4*D*).

Despite the differences in editing efficiency, the editing profiles of WTCasΦ, vCasΦ and nCasΦ were similar. Consistent with previous data for WTCasΦ (19), the vCasΦ and nCasΦ variants also generate deletions primarily around 8- to 10-bp in size (Fig. S6*C*).

### vCasΦ and nCasΦ show higher editing efficiency in transgenic plants

Like in protoplasts, we observed significantly higher editing efficiencies in T1 transgenic plants expressing the vCasΦ and nCasΦ variants compared to WTCasΦ for *AtPDS3* gRNA10 driven by the U6 promoter (Fig. 5*A*). Interestingly, some T1 plants with white sectors were observed from the vCasΦ and nCasΦ expressing T1 populations (Fig. 5*B*), indicating strong editing activity in somatic cells. These white sectors were not observed in transgenic T1 plants expressing wild type CasΦ (Fig. 5*B*). High target gene editing efficiencies were also observed in the T1 plants with vCasΦ and nCasΦ combined with the CmYLCV promoter or UBQ10 promoter driven *AtPDS3* gRNA10 flanked by ribozymes (Fig. 5*C*). Consistent with the higher editing efficiency in the T1 generation, vCasΦ and nCasΦ with *AtPDS3* gRNA10 also generated more albino seedlings in T2 populations compared to the WTCasΦ (Fig. 5*D* and Fig. S7*A*). Transgene free albino seedlings were identified from these T2 populations, showing heritability of the mutations generated by the CasΦ variants (Fig. S7*B*).

**Fig. 5.**
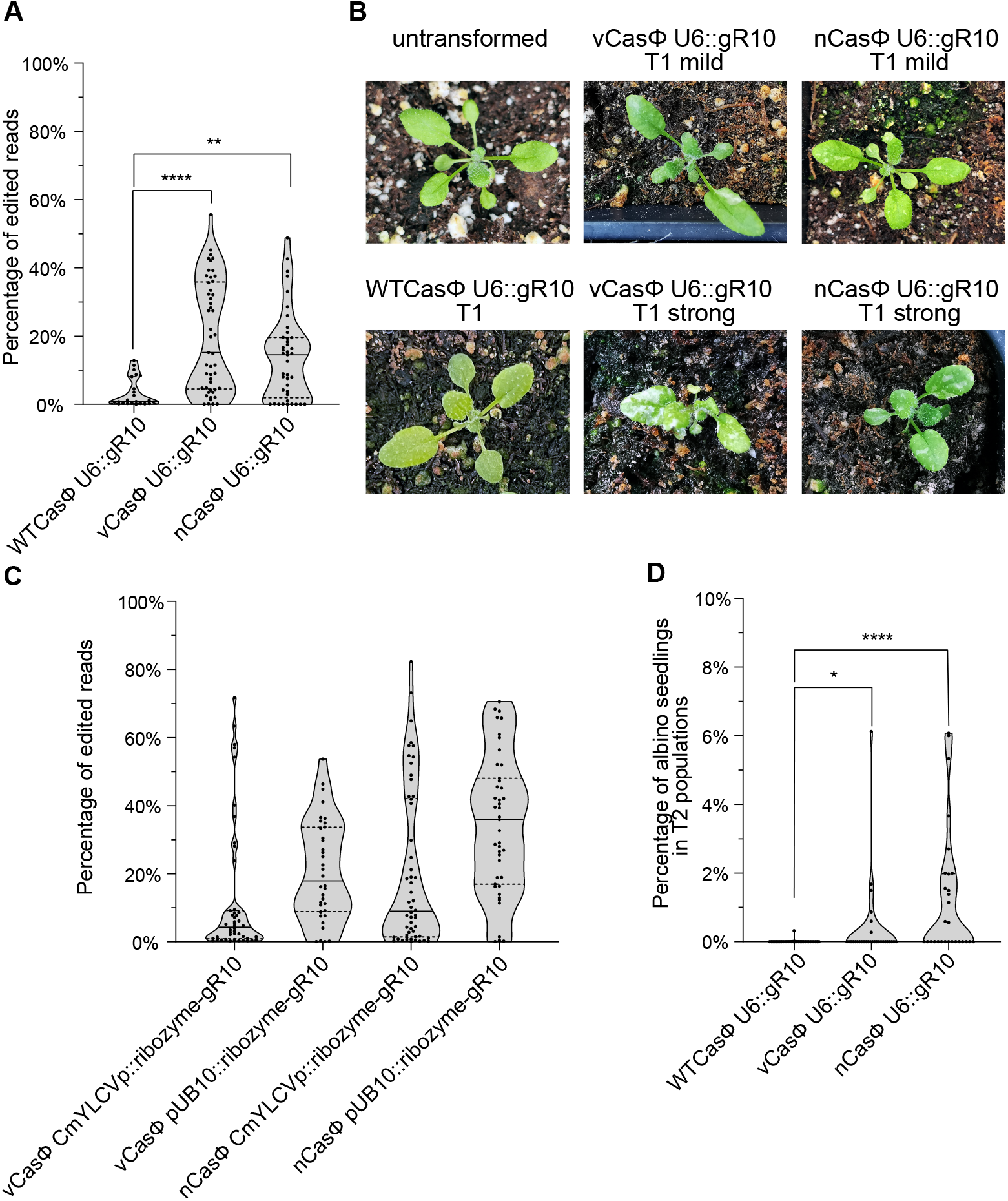
Comparison of editing efficiency of the WTCasΦ with vCasΦ and nCasΦ variants in transgenic plants. (A) Leaf tissue of T1 transgenic plants of indicated constructs in the *rdr6-15* background were harvested for DNA extraction and amplicon sequencing analysis. gR10, *AtPDS3* gRNA10. N=23 for WTCasΦ U6::gR10, n=49 for vCas Φ U6::gR10, and n=41 for nCasΦ U6::gR10. Mann-Whitney test was used to calculate the P value for each comparison indicated. **, 0.001<P<0.01, ****, P<0.0001. (B) A *rdr6-15* plant (untransformed) is shown on the top left, with leaves appearing as uniformly green. Representative T1 transgenic plants in the *rdr6-15* background are shown for WTCasΦ U6::*AtPDS3* gR10 (bottom left), as well as vCasΦ U6::*AtPDS3* gR10 and nCasΦ U6::*AtPDS3* gR10 (top and bottom pictures of representative plants with mild and strong albino patches). (C) Leaf tissue of T1 transgenic plants expressing CasΦ variants with Pol-II promoters and *AtPDS3* gRNA10 flanked by ribozymes in the *rdr6-15* background were harvested for DNA extraction and amplicon sequencing analysis. gR10, *AtPDS3* gRNA10. N=46 for vCasΦ CmYLCVp::ribozyme-gR10, n=36 for vCasΦ pUB10::ribozyme-gR10, n=54 for nCasΦ CmYLCVp::ribozyme-gR10, and n=43 for nCasΦ pUB10::ribozyme-gR10. (D) Total seedling and albino seedling numbers were counted from random T2 populations of WTCASΦ U6::gR10, vCASΦ U6::gR10 and nCASΦ U6::gR10 in the *rdr6-15* mutant background.The percentage of albino seedlings from each T2 population are shown. gR10, *AtPDS3* gRNA10. N=31 for WTCasΦ U6::gR10, n=30 for vCasΦ U6::gR10, and n=29 for nCasΦ U6::gR10. Mann-Whitney test was used to calculate the P value of each comparison indicated. *, 0.01<P<0.05, ****, P<0.0001. In (A), (C) and (D), truncated violin plots and all data points are shown, with median and quartiles indicated by solid and dashed line, respectively.

### Off-target editing by the engineered CasΦ-2 variants vCasΦ and nCasΦ is rare

To evaluate *in vivo* off-target editing frequencies by CasΦ-2 in *Arabidopsis*, we performed whole genome sequencing on transgene free T2 albino seedlings from T1 transgenic plants expressing vCasΦ or nCasΦ *U6::AtPDS3* gRNA10 in the *rdr6-15* mutant background. In T1 transgenic plants with ubiquitous CasΦ expression, unless off-target editing happens at a significant level in sequence contexts similar to the on-target loci, it is difficult to distinguish off-target editing effects from sequence variants generated from spontaneous mutation, PCR amplification errors or sequencing errors. In these transgene free T2 albino seedlings, inherited off-target edits will exist as heterozygous or homozygous mutations, thus making their detection robust by deep sequencing analysis at the whole genome level. Genomic DNA of albino transgene free T2 seedlings from three independent T2 populations of vCasΦ and nCasΦ *U6::AtPDS3* gRNA10 in *rdr6-15*, together with four *rdr6-15* control seedlings, were sequenced to between 116-fold and 613-fold genome coverage, with >99% of genome covered by mapped reads (Fig. 6*A* and Table S2). Variations relative to the reference Col-0 TAIR10 genome sequence were detected by GATK and Strelka2 (35, 36), and similar to previous observations (37–42), a large number of SNP and indels were detected (Table S2). However, the number of variants detected in the *rdr6-15* background plants relative to the Col reference genome were similar to that in the CasΦ-2 edited plants, suggesting that most of the variants detected are variations which already existed in the *rdr6-15* mutant background. We therefore filtered out all variants in the CasΦ-2 edited plants that were also present in the *rdr6-15* background. In addition, to select for heterozygous or homozygous mutations, variants with ratios of reference allele reads over variant allele reads larger than 3 (Ref/Alt>3) were discarded (Fig. 6*A*). Between 66 and 203 variants remained within the six albino seedlings tested after these filtering steps (Table S2). On-target site mutations (*AtPDS3* gRNA10 target region) were reliably detected with this pipeline for all six albino seedlings, with two representative plants shown in Fig. 6*B* left panel. To test if other sequence variants might be due to CasΦ editing, we utilized Cas-OFFinder (43) to predict potential off-target sites with a TBN PAM sequence, allowing up to 4 base pair mismatches and 2 base pairs bulges relative to the *AtPDS3* gRNA10 spacer sequence. We observed no overlap of the predicted off-target sites and the sequence variants detected in the albino seedlings, suggesting that CasΦ-2 editing is highly specific (Table S2). An example of detailed reads at a predicted off-target site is shown in Fig. 6*B* right panel and Fig. S8*A* right panel.

**Fig. 6.**
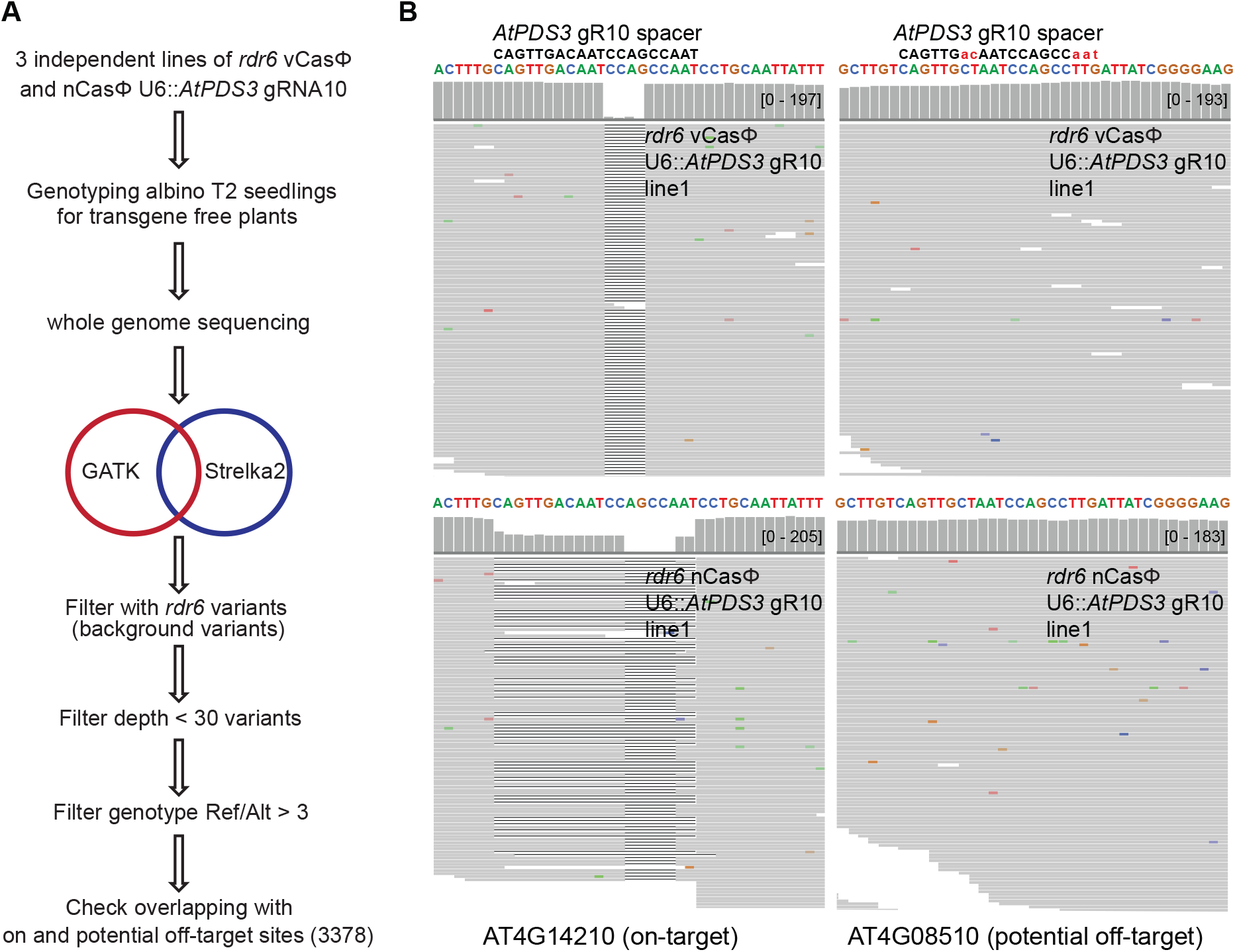
Evaluation of *in vivo* off-target editing by the vCasΦ and nCasΦ variants. (A) Schematic of the workflow to detect potential off-target editing by the vCasΦ and nCasΦ variants through whole genome sequencing. GATK and Strelka2 were used to identify sequence variants compared to the TAIR10 reference genome sequence. Ref/Alt, the ratio of reads with reference sequence over reads with variant sequence. (B) Screenshots of aligned reads and coverage at the *AtPDS3* (*AT4G14210*) gRNA10 target region (left) and a potential off-target site (*AT4G08510*). Top screenshots, from vCasΦ U6::*AtPDS3* gRNA10 line 1. Bottom screenshots, from nCasΦ U6::*AtPDS3* gRNA10 line 1. Capitalized and colored sequences are the reference genomic sequences at these two loci. *AtPDS3* gRNA10 spacer sequence is shown in black letters with uncapitalized red letters showing the mismatched nucleotides between the *AtPDS3* gRNA10 spacer and the potential off-target site.

We also manually inspected each of the sequence variants detected genome wide, which indicated that most of them are located within long stretches of repeated single or di-nucleotides, and reads with similar variants were readily detected in the *rdr6-15* plants, suggesting that most of the detected variants are likely due to imprecise PCR amplification, sequencing reaction or mapping outcomes. After removing these sequence variants at simple repeats, between 6 and 18 high-confidence variants remained in the six albino plants, with more SNP than indels (Fig. S8*B*). However, there was no overlap of the variant loci except for a single SNP between two albino seedlings (Table S3). These data suggest that the high-confidence variants we detected are rare and likely due to spontaneous mutation events, meaning that CasΦ-2 shows little or no off-target activity.

### CasΦ-2 is able to perform gene editing in maize protoplasts

To test gene editing by CasΦ-2 in maize, an agronomically important monocot species with a more complex genome than *Arabidopsis*, eleven guide RNAs were designed targeting the maize *Lc* gene (Table S4). RNPs of WTCasΦ, vCasΦ and nCasΦ with *Lc* gRNAs were transfected into maize protoplasts. By amplicon sequencing, editing of target loci by *Lc* gRNA3 to gRNA7 were detected (Fig. S9). Consistent with results in *Arabidopsis*, vCasΦ and nCasΦ yielded much higher target editing efficiency compared to the WTCasΦ, with up to about 7% editing for the best guide RNA. These data suggest that CasΦ systems might be useful in many different plant species.

## Discussion

In this study, we explored the use of CasΦ-2 for plant genome engineering. We showed that heritable gene edits could be generated in *Arabidopsis* using WTCasΦ at a low frequency. We also showed that the vCasΦ and nCasΦ protein variants exhibited much higher editing efficiency than WTCasΦ in *Arabidopsis* and maize protoplasts, and also a higher frequency of heritable edits in transgenic *Arabidopsis* plants. Given that CasΦ-2 was able to edit the dicot plant *Arabidopsis*, the monocot plant maize, as well as mammalian cells (19), it is likely that CasΦ-2 could be utilized in most eukaryotic systems. The wide temperature range of enzymatic activity of CasΦ-2 may also make it useful for plants and other organisms that grow at cooler temperatures. Since Pol-II promoters can be used for CasΦ-2 guide RNA transcription, the vast array of Pol-II promoters could be useful for expression of gRNAs in specific conditions, such as in different tissues, at higher levels with strong promoters, or in an inducible fashion. CasΦ-2 based editing can also be applied in both RNP and plasmid expression formats. We also found that CasΦ-2 editing was highly specific, with no convincing off-target editing detected by genome wide analysis.

We observed that transgene silencing limited the editing efficiency of CasΦ-2, because editing frequencies were higher when utilizing the *rdr6* mutant. This suggests that the level of expression of either the CasΦ-2, its gRNA precursors, or both, are limiting for high efficiency editing. This also suggests that improvements in transgene designs are likely to lead to more efficient editing. By comparing editing efficiencies at the *FWA* locus in protoplasts derived from the wild type methylated strain or the *fwa* unmethylated epiallele strain, we also found that methylated compact chromatin negatively affected editing by CasΦ-2, presumably because the more highly compact chromatin limits access of CasΦ-2 to the DNA. Fusion of CasΦ-2 protein to chromatin remodelers, as previously reported for the Cas9 protein (44), or other methods, could facilitate successfully editing at difficult target sites, or might increase editing efficiency at all sites.

We observed that more than half of the RNAs we designed yielded no detectable editing events by amplicon sequencing, and only a few of the guide RNAs lead to efficient editing. Similarly, strong differences in editing efficiencies between different guide RNAs were observed when CasΦ-2 was used to target an EGFP gene in HEK293 cells (19). In addition, variable activities were observed with *in vitro* substrate DNA digestion assays with CasΦ-2 RNPs reconstituted with different guide RNAs (Fig. S4). However, with the best performing guide RNAs, reasonable editing efficiencies by CasΦ-2 and its variants were observed. In mammalian cells, more than 30% of cells lost EGFP signal due to the editing of the WTCasΦ-2 with gRNA8 (19). In *Arabidopsis*, with the CasΦ-2 protein variants and *AtPDS3* gRNA10, 10% to 30% editing efficiency was observed in protoplasts (Fig. 4*D*) and 40-80% of somatic editing efficiency was observed in a multiple T1 transgenic plants (as indicated by the upper quartile in Fig. 5*C*). However, in the future it will be important to systematically define gRNA design principles to increase these frequencies.

Amplicon sequencing and *in vivo* editing results indicated that CasΦ-2 edits consist of small deletions of roughly nine base pairs, consistent with biochemical data showing that CasΦ-2 produces an 8 to 12 nucleotide staggered cut (19). CasΦ-2 could therefore be useful for making in-frame deletions to create partial loss of function alleles, or to remove small sections of specific domains of proteins. Indeed, one of the heritable edits found in this study was a two amino acid deletion resulting in an in-frame deletion of the coding region of the *AtPDS3* gene. The ability to make small deletions may also be useful for promoter analysis by removing individual transcription factor binding sites, or by promoter deletion scanning by tiling gRNAs across a promoter.

In summary, this study demonstrates that the compact CasΦ editing system can be used for gene editing in plants and can generate heritable mutations with high specificity. Compared to the Cas9 and Cas12a systems, which has been optimized with community efforts for years, CasΦ-2 is still less efficient in overall editing efficiency. However, with well-performing guide RNAs, a reasonable level of editing efficiency can be obtained with the CasΦ system. In addition, the vCasΦ and nCasΦ variants demonstrated the effectiveness of protein engineering to potentially improve the editing efficiency further in the future. With its hypercompact protein size, wide working temperature range and T-rich minimal PAM, CasΦ should be a useful supplement to the plant gene editing toolbox, and should be useful in both basic research and agricultural biotechnology.

## Materials and Methods

### Plant materials and growth condition

To grow *Arabidopsis* plants for protoplast preparation, Col-0 and *fwa-4* epi-mutant plants were grown under a 12-hour light/12-hour dark photoperiod and with low light condition for 3-4 weeks. To grow maize plants for protoplast isolation, B73 seeds were soaked in water overnight and planted in half peat moss and half vermiculite. After three days of light incubation, emerged seedlings were transferred to dark until the 2^nd^ leaf was 10-15 cm long.

Agrobacterium mediated *Arabidopsis* plant transformation was performed as previously described (45) and transgenic T1 plants were screened with half MS plates with 40 μg/ml hygromycin B. For 28 °C treatment of T1 plants, including the T1 plants in Fig. S1*B* (version2 construct *AtPDS3* gRNA10), stratified seeds on half MS plates with hygromycin B were incubated at 28 °C and resistant T1 plants were transferred to soil after about a week, and put back to 28 °C for a total of two weeks incubation at 28 °C. T1 plants were then moved to a greenhouse (23 °C) for the rest of the life cycle. To support the growth of albino seedlings in T2 generation, 3% sucrose was supplemented to half MS plates when needed.

### Protein purification

WTCasΦ, vCasΦ and nCasΦ proteins with NLS were purified as previously described (19, 22).

### RNP reconstitution

Guide RNAs were synthesized as 25 nucleotide repeat + 20 nucleotide spacer as shown in Table S4. Lyophilized RNA was dissolved by adding DEPC-treated H_2_O to a concentration of 0.5 mM. The dissolved RNA was incubated at 65 °C for 3 min, then cooled down to room temperature. For RNP reconstitution, heated and cooled RNA was added to 2xCleavage Buffer (2xCB buffer, 20 mM Hepes-Na, 300 mM KCl, 10 mM MgCl_2_, 20% glycerol, 1 mM TCEP, pH 7.5) to a final concentration of 5 μM and vortexed to mix. Then, WTCasΦ, vCasΦ or nCasΦ proteins were added to a final concentration of 4 μM and mixed by pipetting. This solution was then incubated at room temperature for 30 min. The resulting solution contains 4 μM of RNP in 2xCB buffer.

### Plasmids used in this study

Plasmids generated in this study are listed in Table S5. The CASΦ-2xSV40NLS-2xFLAG coding sequence (without IV2 intron) was *Arabidopsis* codon optimized and synthesized by IDT. The HBT-pcoCAS9 vector (addgene52254) backbone with the FLAG and SV40 NLS (for version1) or without the FLAG and SV40NLS (for version 2) were amplified and assembled by TAKARA in-fusion HD cloning kit (cat639650) with the synthesized CasΦ sequence amplified as two fragments as well as the IV2 intron sequence amplified from the HBT-pcoCAS9 vector. pCAMBIA1300-pYAO-cas9-MCS vector (46) was digested by KpnI and EcoRI to remove the pYAO-cas9 cassette. Then, pcoCasΦ with IV2 intron, NLS and FLAG tag coding sequence were amplified from HBT_pcoCASphi_version1 and version2 plasmids and cloned into the digested vector together with PCR amplified pUB10 and Rbcs E9 terminator by TAKARA in-fusion HD cloning kit (cat639650). Subsequently, various guide RNA cassettes were cloned into the SpeI site of the obtained vectors by TAKARA in-fusion reaction or by restriction digestion and ligation of PCR amplified or IDT synthesized DNA fragments. Construction of the nCasΦ and vCasΦ binary vectors were performed by first digesting the pC1300_pUB10_pcoCASphi_E9t_MCS_version2 vector with KpnI to remove pUB10 promoter and the first part of CasΦ coding sequence. Then, pUB10 promoter and CasΦ sequences containing the desired mutations were PCR amplified as two fragments with desired mutations added by overlapping primers, and cloned back into the digested vector by an in-fusion reaction. Primers used for cloning and synthesized DNA fragments used for cloning are listed in Table S6. The control plasmid HBT-sGFP for protoplast transfection was previously described (47) and obtained from ABRC (stock CD3-911).

### Protoplast isolation and transfection

*Arabidopsis* mesophyll protoplast isolation was performed as previously described (47). For RNP transfection into *Arabidopsis* protoplasts, 26 μl of 4 μM RNP was first added to a round bottom 2 ml tube, followed by 200 μl of protoplasts (2×10^5^ cells/ml). Then, 2 μl of 5 μg/μl salmon sperm DNA was added and mixed gently by tapping the tube 3-4 times. Finally, 228 μl of fresh, sterile and RNase free PEG-CaCl_2_ solution (47) was added to the protoplast-plasmid mixture and mixed well by gently tapping the tube. The protoplasts with PEG solution were incubated at room temperature for 10 min, then, 880 μl of W5 solution (47) was added and mixed with the protoplasts by inverting the tube 2-3 times to stop the transfection. Protoplasts were harvested by centrifuging the tubes at 100 rcf for 2 min and resuspended in 1 ml of WI solution. They were then plated in 6-well plates pre-coated with 5% calf serum. These 6-well plates were then incubated at room temperature for 48 hours.

For plasmid transfections into *Arabidopsis* protoplasts, the concentrations of plasmids were determined by nanodrop. Then 20-50 μg of plasmids (plasmid amounts were the same within each experiment) were added to the bottom of each transfection tube, and the volume of the plasmids was supplemented with H_2_O to reach 20 μl. 200 μl of protoplasts were added followed by 220 μl of fresh and sterile PEG-CaCl_2_ solution. The mixture was mixed well by gently tapping tubes and incubated at room temperature for 10 min. 880 μl of W5 solution was added and mixed with the protoplasts by inverting the tube 2-3 times to stop the transfection. Protoplasts were harvested by centrifuging the tubes at 100 rcf for 2 min and resuspended in 1 ml of WI solution. They were then plated in 6-well plates pre-coated with 5% calf serum. These 6-well plates were then incubated at room temperature or at 28 °C for 48 hours.

For maize protoplast isolation, middle 8-10 cm of the 2^nd^ leaf from etiolated maize seedlings were sliced into thin strips (about 0.5 mm wide) and immersed in enzyme solution (0.6 M mannitol, 10 mM MES pH 5.7, 1.5% cellulase R10, 0.3% macerozyme R10, 1 mM CaCl_2_, 5 mM BME, 0.1% BSA). Then the enzyme solution with leaf stripes was put under vacuum for half an hour then placed in dark for 2.5 hours. The solution was then gently swirled to release the protoplasts and filtered through a 40 μm cell strainer. The cells were collected by centrifugation at 150 g for 5 min, and resuspended in 10 ml washing solution (0.6 M mannitol, 4 mM MES pH 5.7, 20 mM KCl). Then the cells were rested on ice for at least half an hour. After resting, cell pellets were resuspended with washing solution to 2×10^^5^ cells/ml in cold washing buffer. For RNP transfection into maize protoplasts, 13 μl 4 μM RNP and 1μl 5 μg/μl salmon sperm DNA were added to 100 μl protoplasts. Then 114 μl PEG solution (40% PEG 4000, 0.2 M mannitol, 0.1 M CaCl_2_) were added and mixed by tapping. After 10 min of incubation, transfections were stopped by adding 880 μl washing solution and gently inverting several times. Transfected protoplasts were collected by 3 min of centrifugation at 150 g and resuspended in 1 ml incubation buffer (48). Then, the protoplasts were plated in 6-well plates pre-coated with 5% calf serum. These 6-well plates were then incubated at room temperature for 48 hours.

### Amplicon sequencing

DNA was extracted from protoplast samples and leaves of transgenic plants with Qiagen DNeasy plant mini kit (Qiagen 69106). The amplicon was obtained using two rounds of PCR. Amplification primers for the first round of PCR were designed to have the 3’ sequence of the primers flanking a 200-300 bp fragment of the genomic area targeted by the guide RNA of interest. Primers for the first round of amplification are listed in Table S6. The 5’ part of the primer contained a sequence which will be bound by common sequencing primers. After 25 cycles of the first round of PCR amplification, the reaction was purified using 1x Ampure XP beads (Beckman Coulter A63881).

The eluate was used as template for the second round of PCR amplification of 12 cycles. The second round of PCR was designed so that indexes were added to each sample. The samples were then purified using 0.8x Ampure XP beads. Part of the purified libraries were run on a 2% agarose gel to check for size and absence of primer dimer (fragments below 200 bp considered as primer dimer). Then amplicons were sent for next generation sequencing.

### Amplicon sequencing result analysis

Reads were first quality and adaptor trimmed using *Trim Galore* and then mapped to the target genomic region by the BWA aligner (v0.7.17, BWA-MEM algorithm). Sorted and indexed bam files were used as input files for further analysis by the CrispRvariants R package (v1.14.0). Each mutation pattern with corresponding read counts was exported by the CrispRvariants R package. After assessing all control samples, a criterion to classify reads as edited reads was established: only reads with a >= 3 bp deletion or insertion (indel, mainly as deletions) of the same pattern (indels of same size starting at the same location) with >= 100 read counts from a sample were counted as edited reads. This criterion is established due to the observation of 1 bp indels and occasionally 2 bp indels with read numbers >100 in control samples. Also, larger indels that occur at very low frequencies (much lower than 100 reads) were observed in control samples. These observations indicate that occasional PCR inaccuracy and low-quality sequencing in a small fraction of reads can result in the indel patterns with corresponding read number ranges as stated above in control samples with the typical sequencing depth in our experiments (1-5 million reads/sample). By employing such stringent criteria, it is believed that the editing signals counted are true signals indicating editing events. Additionally, for *FWA* gRNA5 and gRNA6 targeted regions, there are long stretches of adenines near these target regions. Due to the high error rate of polymerases amplifying long stretches of adenines, reads with indels only within these stretches of adenines were not counted as true deletions. For amplicon sequencing result analysis of maize protoplast, the criterion to classify reads as edited reads was established: only reads with a >= 3bp deletion or insertion (indel, mainly as deletions) of the same pattern (indels of same size starting at the same location) with >= 10 read counts from a sample were counted as edited reads. The adjustment of the criterion was based on the fact that these amplicon sequencing was of less depth (performed by the iSeq 100) compared to the amplicon libraries of *Arabidopsis* protoplasts.

### Off-target analysis

DNA from single *Arabidopsis* seedlings was extracted with the Qiagen DNeasy plant mini kit and sheared to 300 bp size with a Covaris. Library preparation was performed with Tecan Ovation Ultralow V2 DNA-seq kit. For variant calling, whole genome sequencing reads were aligned to the TAIR10 reference genome using BWA mem (v0.7.17) (49) with default parameters. GATK (4.2.0.0) (35) MarkDuplicatesSpark was used to remove PCR duplicate reads. Then GATK HaplotypeCaller was used to call raw variants. Raw SNPs were filtered with QD < 2.0, FS > 60.0, MQ < 40.0, and SOR > 4.0. Raw InDels were filtered with QD < 2.0, FS > 200.0, and SOR > 10.0 and used for base quality score recalibration. The recalibrated bam file was further applied to GATK and Strelka (v2.9.2) (36) for SNPs/InDel calling. Only SNPs/InDels called by both GATK and Strelka were used for further analysis. The intersection of GATK and Strelka SNPs/InDels were filtered by removing identical SNPs/InDels in the *rdr6-15* background by BedTools (v2.26.0) (50). Variants with coverage lower than 30 reads were removed. Variant loci at which the ratio of the reads with the WT allele over the reads with variant allele larger than three were removed. For heterozygous alleles, at a coverage level of 30 reads, the chance of observing WT reads /mutation allele reads >3 is 0.26% (binomial distribution, one-tailed P=0.0026).

## Data deposition

All high-throughput sequencing data generated in this study are accessible at the National Center for Biotechnology information Gene Expression Omnibus via series accession GSE206798. (also weblink here https://www.ncbi.nlm.nih.gov/geo/query/acc.cgi?acc=GSE206798)

## Acknowledgments

We thank Suhua Feng and Mahnaz Akhavan for support with high-throughput sequencing at the UCLA Broad Stem Cell Research Center BioSequencing Core Facility. S.E.J and J.A.D. are Investigators of the Howard Hughes Medical Institute. P.P. receives funding from the European Regional Development Fund under grant agreement number 01.2.2-CPVA-V-716-01-0001 with the Central Project Management Agency (CPVA), Lithuania, and from the Research Council of Lithuania (LMTLT) under grant agreement number S-MIP-22-10.

## Author contributions

Z.L. and S.E.J. designed the experiments. Z.L., Z.W. and J.A. performed the experiments. Z.L. and Z.Z. analyzed the results. P.P. purified proteins for RNP assay. B.A.S. helped design of guide RNAs. P.P., B.A.S. and J.A.D. contributed in discussions and designing of experiments. Z.L. and S.E.J. wrote the paper with input from the other authors.

