## supplement figure and table for "Genome editing in plants using the compact editor CasΦ"

**This PDF file includes:**

Supporting text
Figures S1 to S9
Tables S1 to S6

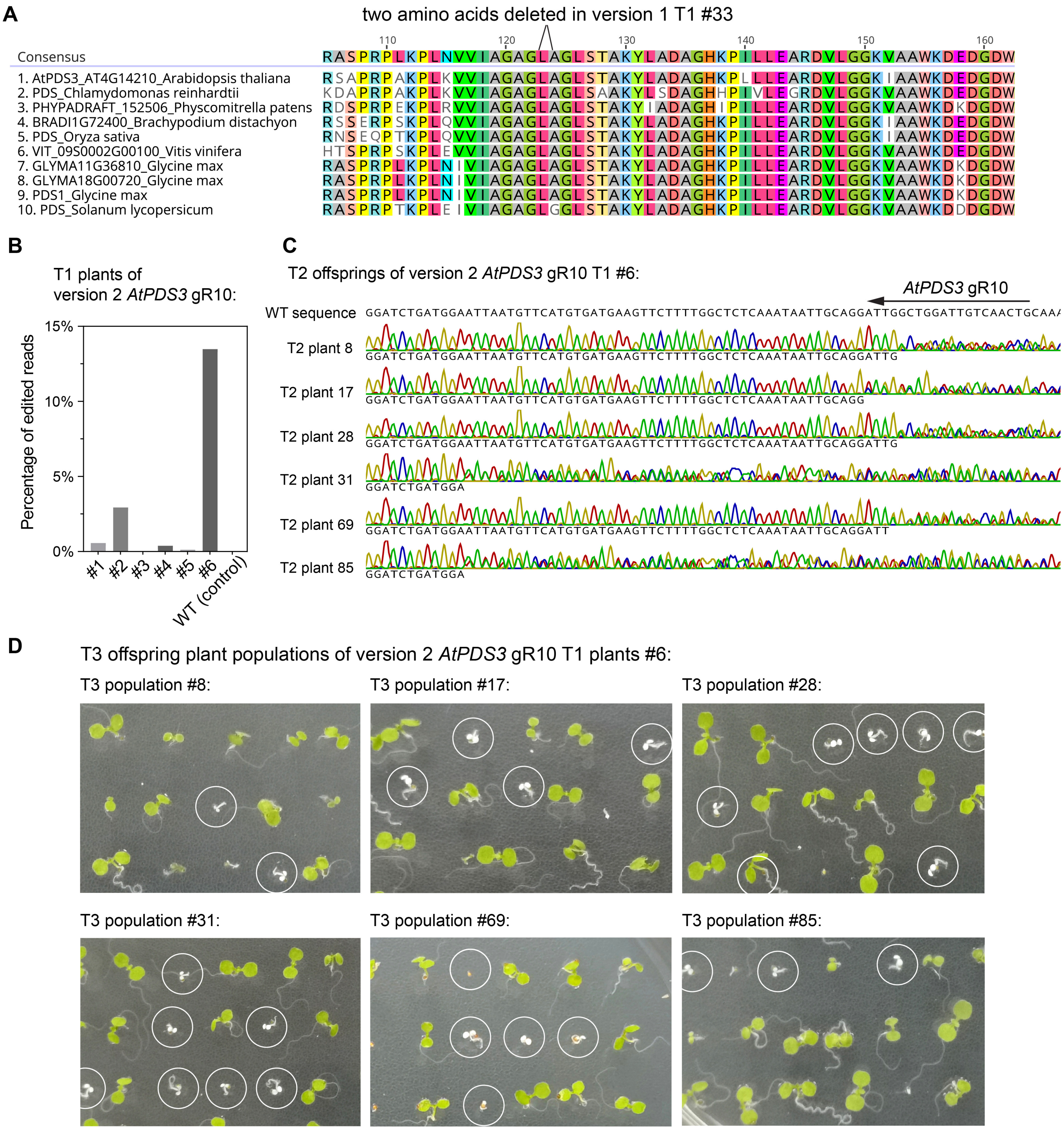

**Fig. S1.** CasΦ-2 mediated editing of the *AtPDS3* gene with version 1 and version 2 constructs. (A) AtPDS3 homolog protein sequences from different species were aligned with Clustal Omega by the Geneious software, with the two amino acids deleted in version 1 construct T1 plant #33 labeled. (B) Amplicon sequencing results of T1 plant leaves of version 2 *AtPDS3* gRNA10 construct. (C) Sanger sequencing results of the *AtPDS3* gRNA10 target region of six out of 96 total seedlings from the T2 population of version 2 *AtPDS3* gR10 T1 plant #6, showing that they are heterozygous for mutation in this region. (D) Seedlings of six T3 offspring plant populations of version 2 *AtPDS3* gRNA10 T1 plant #6, corresponding to the six T2 plant indicated in (B), with albino seedlings circled.

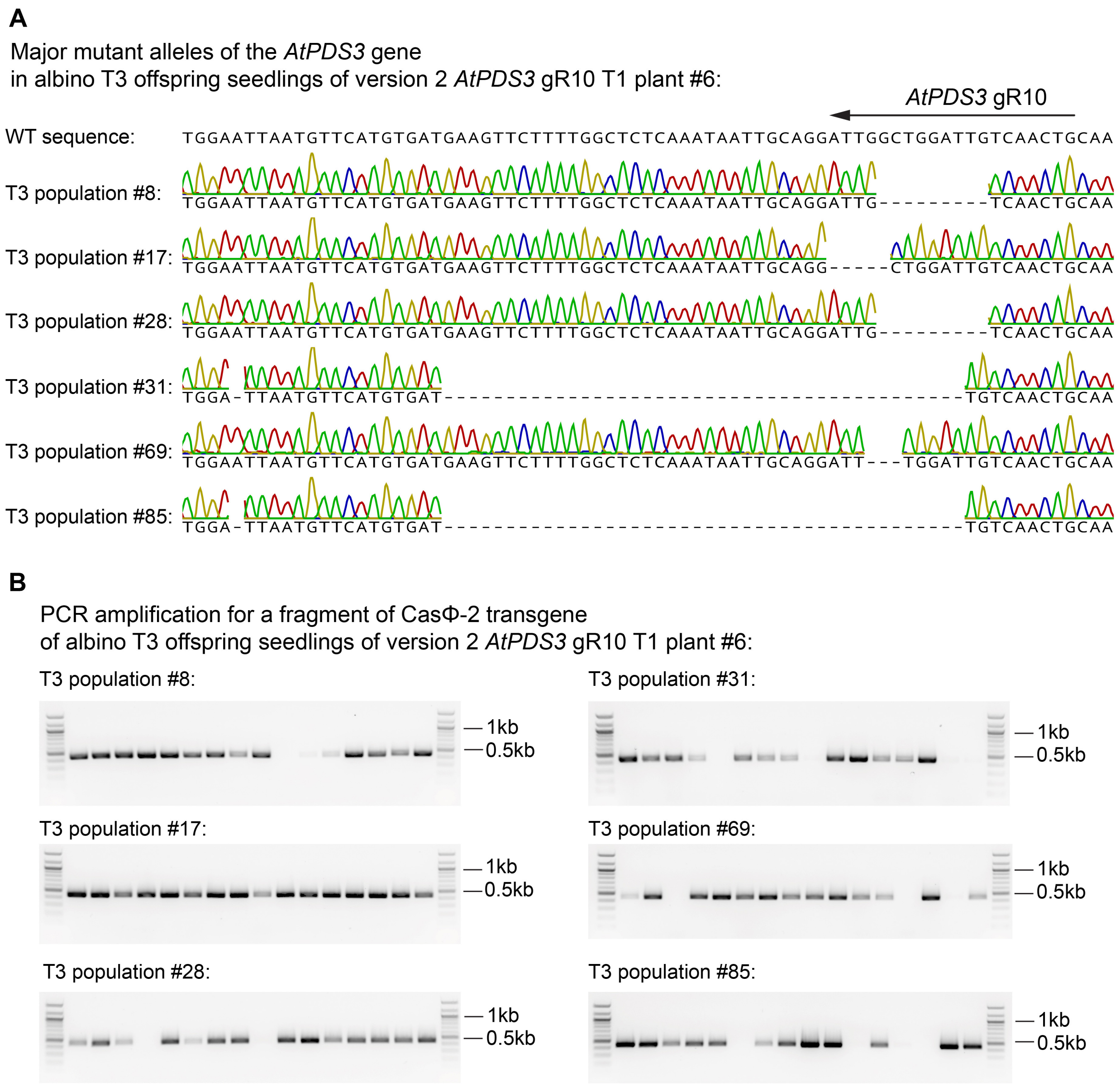

**Fig. S2.** Characterization of the editing of the *AtPDS3* gene by CasΦ-2 with the version 2 construct in the T3 generation. (A) multiple albino seedlings from T3 populations corresponding to the T2 plants in Fig. S1*C* were sanger sequenced and the major mutant alleles are displayed. (B) PCR amplification of the DNA of 16 randomly selected albino seedlings from the T3 populations in (A) for a fragment of the CasΦ-2 transgene.

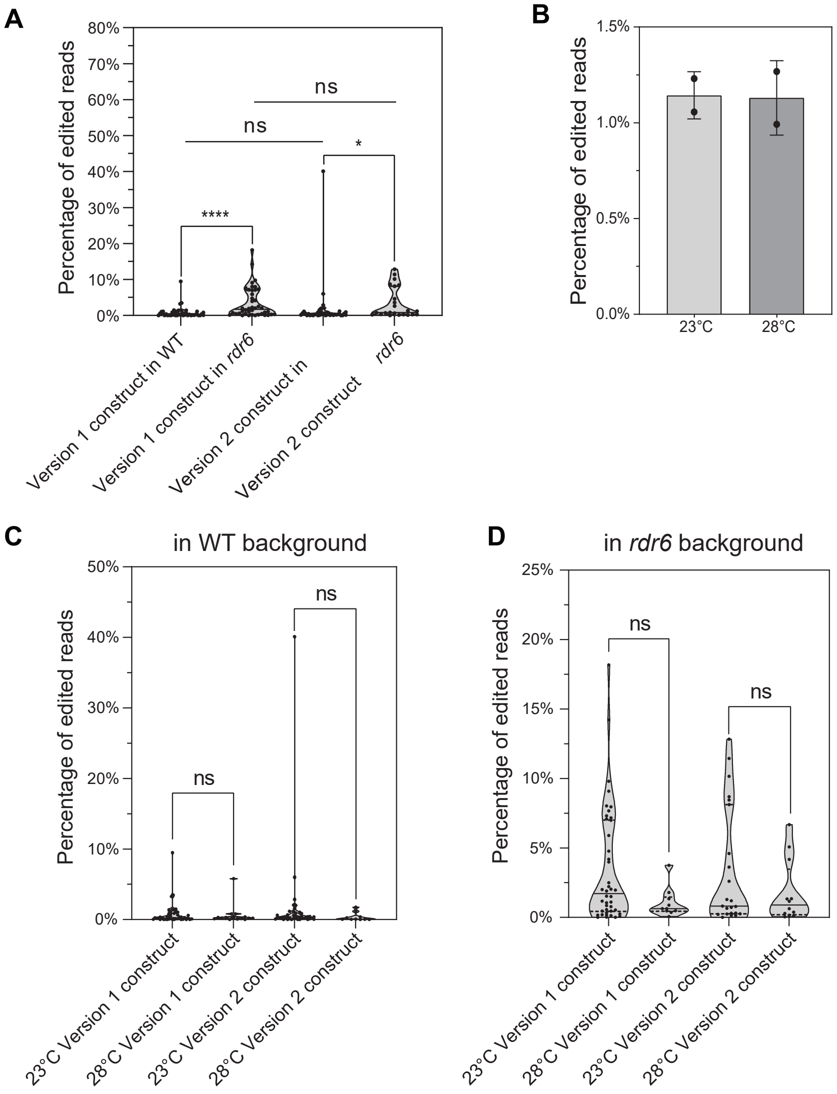

**Fig. S3.** The impact of transgene silencing and temperature on target gene editing efficiency by CasΦ-2. (A) Version 1 and version 2 constructs with *AtPDS3* gRNA10 were transformed into Col-0 (WT) and *rdr6-15* (*rdr6*) backgrounds. Editing efficiencies in T1 leaves determined by amplicon sequencing were plotted. N=43 for version 1 construct in WT background, n=42 for version 1 construct in *rdr6* background, n=41 for version 2 construct in WT background and n=23 for version 2 construct in *rdr6* background. (B) The Col-0 protoplasts were transfected with version 2 construct with *AtPDS3* gRNA10 and incubated at 23°C and 28°C. Two replicate transfections were performed for each temperature and the editing efficiency of each individual transfection, as well as the mean and standard deviation of the two replicates were plotted. (C) and (D), version 1 and version 2 constructs with *AtPDS3* gRNA10 were transformed into WT background (C) and the *rdr6* mutant background (D). T1 plants were incubated constantly at 23°C (23°C sets) or initially at 28°C for 2 weeks then at 23°C (28°C sets). Editing efficiencies in T1 leaves were plotted. In the WT background (C), n=43 for the set of 23°C version 1 construct, n=15 for the set of 28°C version 1 construct, n=41 for the set of 23°C version 2 construct and n=10 for the set of 28°C version 2 construct. In the *rdr6* background (D), n=42 for the set of 23°C version 1 construct, n=10 for the set of 28°C version 1 construct, n=23 for the set of 23°C version 2 construct and n=12 for the set of 28°C version 2 construct. In (A), (C) and (D), truncated violin plots and all data points are shown, with median and quartiles indicated by solid and dashed line, respectively. Mann-Whitney test was used to calculated the P value for each comparison indicated. ns, non-significant, P>0.05; *, 0.01<P<0.05; ****, P<0.0001.

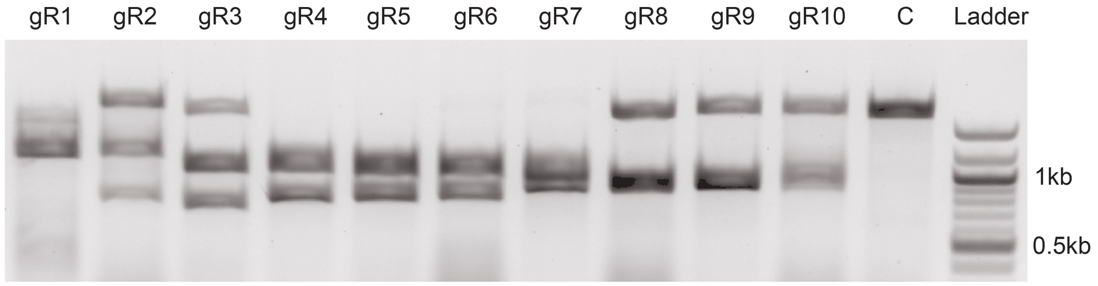

**Fig. S4.** *In vitro* cleavage of PCR amplified *FWA* gene fragment by CasΦ -2 RNP with *FWA* gRNA1 to gRNA10. A1.57kb *FWA* gene fragment spanning all gRNA target regions was amplified by PCR and gel purified. The *FWA* gene f ragment wa s incubated with Ca sΦ-2 RNPs containing gRNA1 to gRNA10 (gR1 to gR10) and a scrambled gRNA control (C) at 37°C for 1 hour. Reactions were stopped by adding EDTA and digestion o f C asΦ-2 protein with proteinase K. 2% agarose gels were used to visualize the cleavage products along with a DNA ladder for sizing.

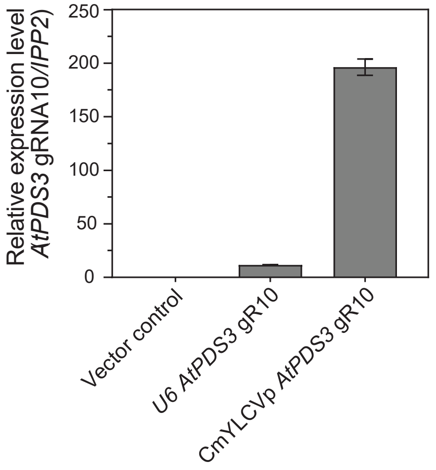

**Fig. S5.** Comparison of the level of the *AtPDS3* gRNA10 driven by the *U6* and the *CmYLCV* promoter in protoplasts. Real-time quantitative PCR was used to measure the level of *AtPDS3* gRNA10 expression level in protoplasts transfected with the same amounts of the version 2 U6::*AtPDS3* gRNA10 plasmid and the version 2 CmYLCVp::*AtPDS3* gRNA10 plasmid. Protoplasts transfected with the version 2 U6::*AtPDS3* gRNA8 plasmid was used as the vector control to evaluate basal noise level of the primer pair used for the *AtPDS3* gRNA10 amplification. The *IPP2* gene was used as a reference gene for normalization. Three technical replicates were performed. Mean and standard error of the relative quantity calculated by the Bio-Rad CFX software are plotted.

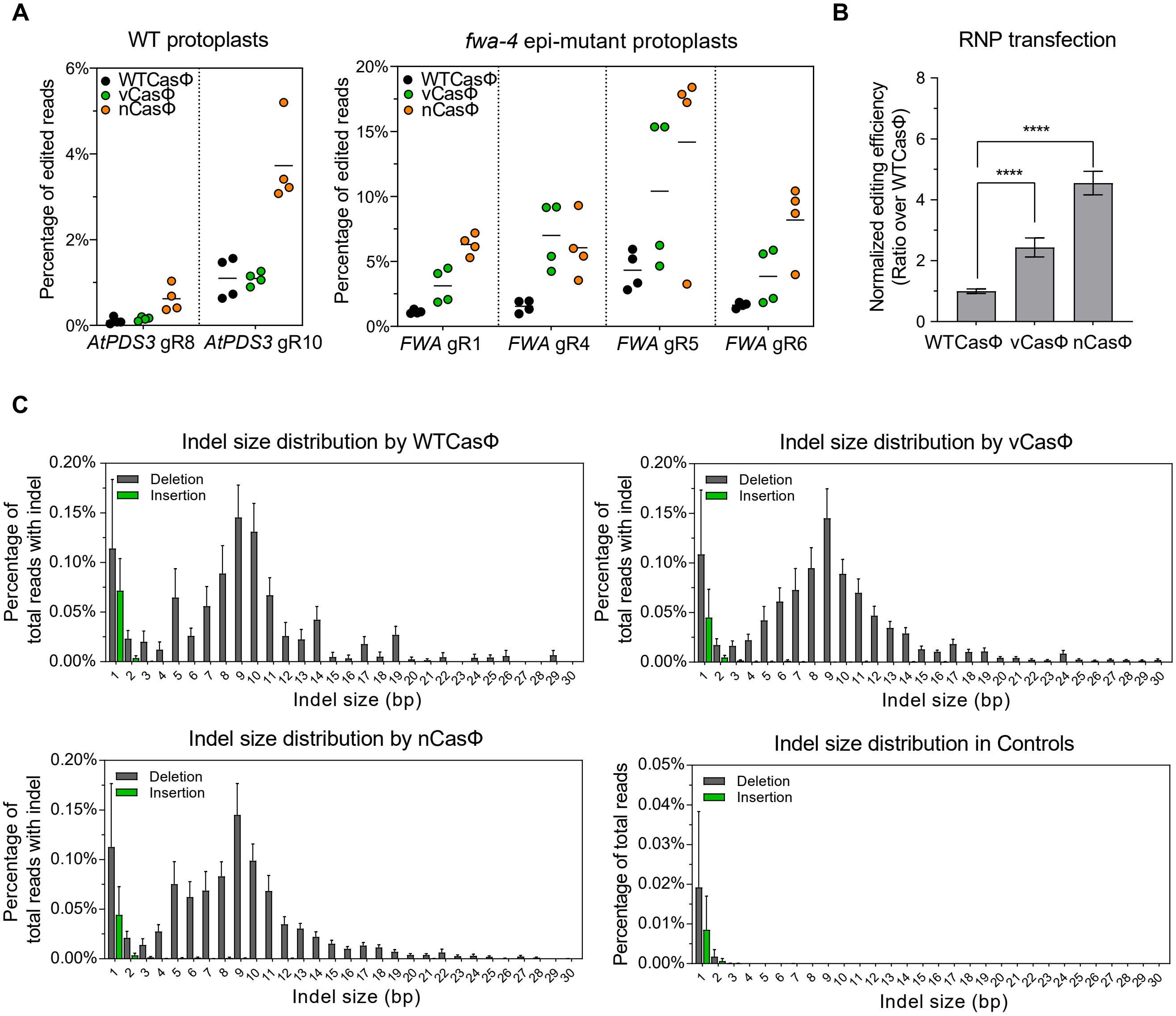

**Fig. S6.** Comparison of editing efficiency and indel size distribution profile by the vCasΦ and nCasΦ variants and WTCasΦ in protoplasts. (A) RNPs reconstituted with WTCasΦ, vCasΦ and nCasΦ proteins and guide RNAs as indicated were transfected into protoplasts prepa red from Col-0 plants (WT) (left panel) and from *fwa-4* epi-mutant plants (right panel). Individual replicate values and mean of the four replicates of each test were plotted. (B) Target gene editing efficiencies in (A) were normalized by calculating the ratio of editing efficiencies over that of mean editing efficiency by WTCasΦ for each guide RNA. Mean and standard error of the normalized editing efficiencies for all gRNAs were plotted. Unpaired t-test was used to calculate P value of indicated comparisons. ****, P<0.0001. (C) The indel size distribution was calculated as the percentage of reads of a particular insertion or deletion size, from 1 bp to 30 bp, among all reads with indels for each protoplast transfections in Fig. 4*A*. Mean and standard error of the indel size distributions of all guide RNAs in Fig. 4*A* are plotted. For the control samples (bottom right panel), protoplasts transfected with the HBT-sGFP plasmid were amplified for the target regions of the six guide RNAs used in Fig. 4*A* (six amplicon sequencing for the control samples in total). The indel size distribution was calculated as the percentage of reads of a particular insertion or deletion size among all reads.

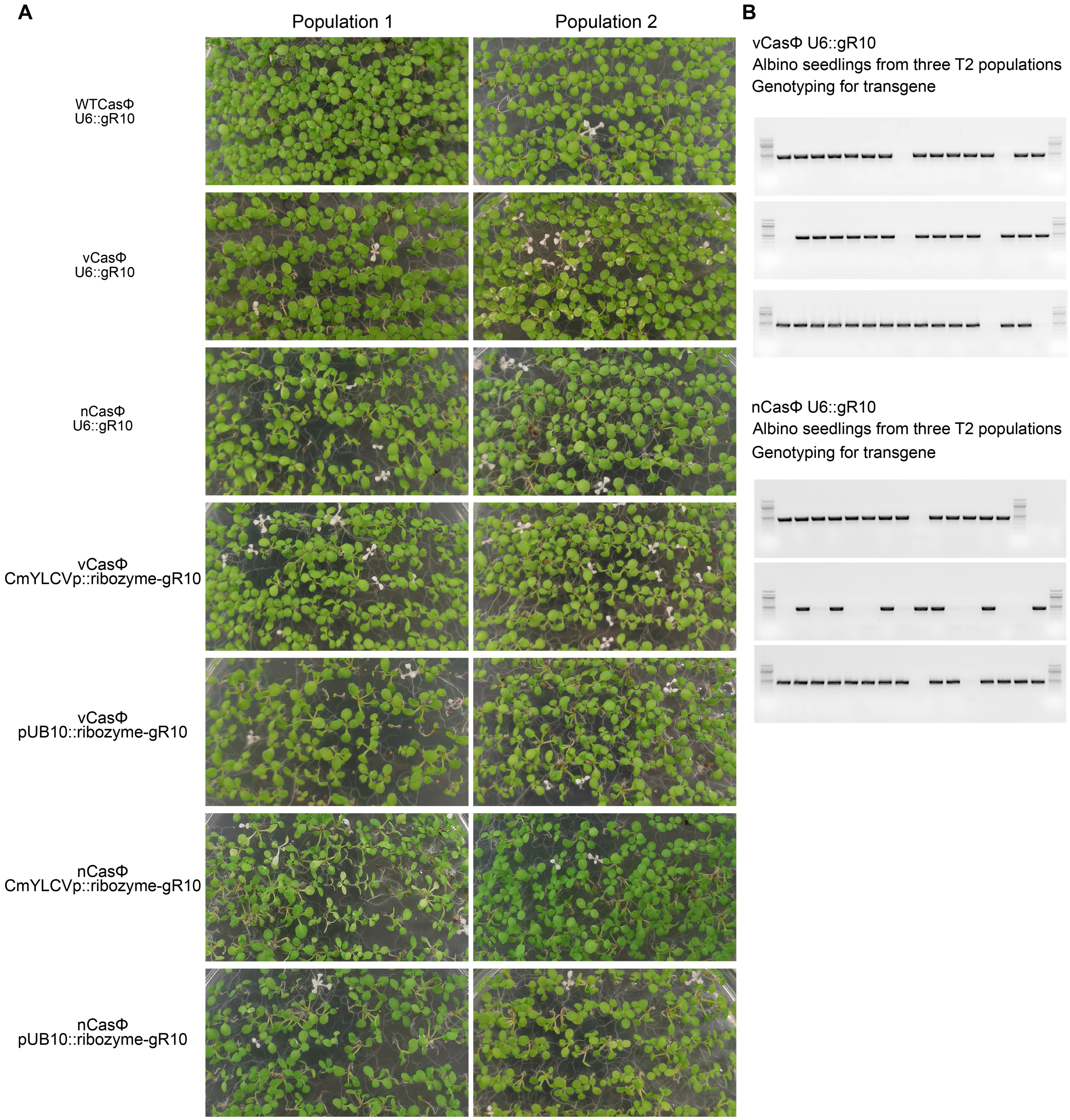

**Fig. S7.** Heritability analysis of the editing of the *AtPDS3* gene by the vCasΦ and nCasΦ variants in the T2 generation. (A) Pictures of two representative T2 populations of transgenic plants of indicated constructs in the *rdr6-15* background. gR10, *AtPDS3* gRNA10. (B) PCR amplification of the DNA of randomly selected albino seedlings from three T2 populations of vCASΦ and nCASΦ U6::*AtPDS3* gR10 in the *rdr6-15* background for a fragment of the CasΦ-2 transgene.

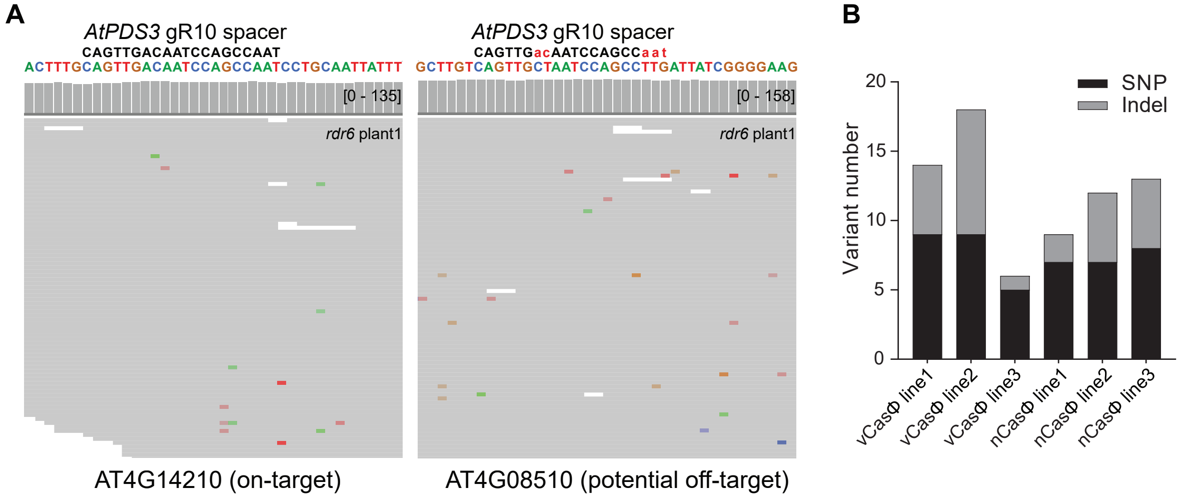

**Fig. S8.** Summary of high confidence variants discovered in the transgene-free albino T2 seedlings. (A) Screenshots of aligned reads and coverage of a control *rdr6-15* plant at the *AtPDS3* (*AT4G14210*) gRNA10 target region (left) and a potential off-target site (*AT4G08510*). Capitalized and colored sequences are the reference genomic sequences at these two loci. *AtPDS3* gRNA10 spacer sequence is shown in black letters with uncapitalized red letters showing the mismatched nucleotides between *AtPDS3* gRNA10 spacer and the potential off-target site. (B) The number of high confidence SNPs and Indels identified genome wide for the sequenced transgene free albino seedlings.

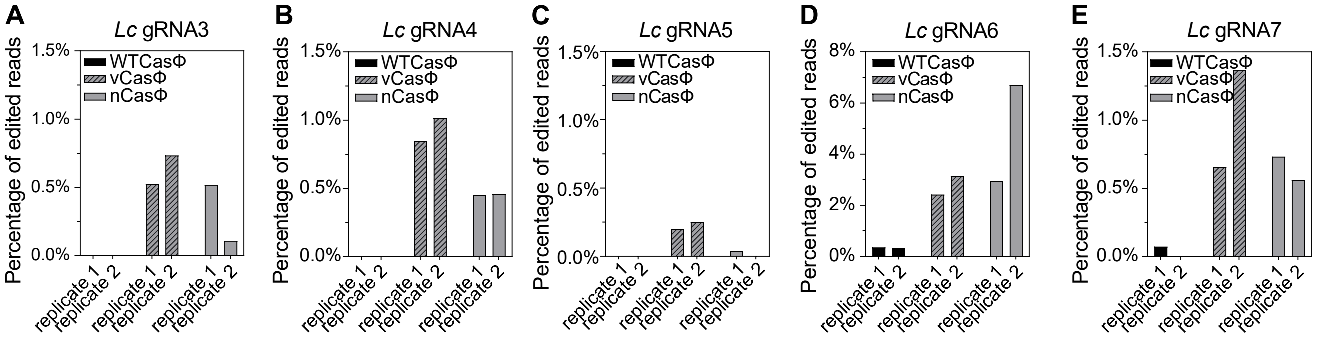

**Fig. S9.** The editing of *Lc* gene by CasΦ in maize protoplasts. (A)-(E) RNPs reconstituted with WTCasΦ, vCasΦ and nCasΦ proteins and 11 guide RNAs targeting the maize Lc gene were transfected into maize protoplasts. Editing of the target region was detected with five out of the 11 guide RNAs tested and editing efficiencies of the two replicate transfections are shown. (A) *Lc* gRNA3. (B) *Lc* gRNA4. (C) *Lc* gRNA5. (D) *Lc* gRNA6. (E) *Lc* gRNA7.

**Table S1.** Detailed editing events detected in version 1 construct with *AtPDS3* gRNA10 in T1 plant # 33.

| tissue | editing event* | supporting reads | total edited reads | total reads | editing efficiency (%) |
|---|---|---|---|---|---|
| flower 1 | -5:6D | 284 | 284 | 3383814 | 0.0083929 |
| leaf 1 | -5:6D | 1656570 | 1660178 | 3671170 | 45.22204093 |
| | -4:6D | 2728 | | | |
| | -6:6D | 657 | | | |
| | -1:5D | 114 | | | |
| | -2:6D | 109 | | | |
| leaf 2 | -5:6D | 1922514 | 1926043 | 3887866 | 49.5398504 |
| | -4:6D | 2701 | | | |
| | -6:6D | 697 | | | |
| | -2:6D | 131 | | | |
| leaf 3 | -5:6D | 425 | 529 | 3616680 | 0.01462667 |
| | -4:7D | 104 | | | |
| leaf 4 | -5:6D | 425481 | 426469 | 3954723 | 10.78378941 |
| | -4:6D | 882 | | | |
| | -6:6D | 106 | | | |

*Editing events are shown as: (position where the editing starts) : (number of nucleotides of) D (deletion) or I (insertion). position 0 is between the 18th and 19th nucleotides of the guide, so that the 18th nucleotide is position -1, the 19th nucleotide is position +1.

**Table S2.** Summary of the analysis of whole genome sequencing data of the *rdr6-15* control and CasΦ transgene free T2 albino seedlings.

| | | | | | | T2 transgene negative plant | | | | | |
|---|---|---|---|---|---|---|---|---|---|---|---|
| | | *rdr6* control | | | | *rdr6 vCasphi U6PDS3gR10* | | | *rdr6 nCasphi U6PDS3gR10* | | |
| | | plant 1 | plant 2 | plant 3 | plant 4 | line1 | line2 | line3 | line1 | line2 | line3 |
| Reads Number | | 104272905 | 244547548 | 57313755 | 46421199 | 150763760 | 87663735 | 136018868 | 117908027 | 126570163 | 154534923 |
| Reads Length | | 150 | 150 | 150 | 150 | 150 | 150 | 150 | 150 | 150 | 150 |
| Reads Type | | PE | PE | PE | PE | PE | PE | PE | PE | PE | PE |
| Coverage | | 261.41 | 613.07 | 143.68 | 116.38 | 377.96 | 219.77 | 340.99 | 295.59 | 317.30 | 387.41 |
| Mapped reads | | 102752061 | 241841799 | 56528541 | 45991324 | 148973332 | 86547235 | 134468420 | 116178427 | 125262252 | 152400339 |
| Reads Mapping Rate | | 98.5415% | 98.8936% | 98.6300% | 99.0740% | 98.8124% | 98.7264% | 98.8601% | 98.5331% | 98.9667% | 98.6187% |
| Duplication Rate | | 18.76% | 20.72% | 10.98% | 9.22% | 22.70% | 17.06% | 19.49% | 17.93% | 20.99% | 19.87% |
| Covered Genome | | 99.8307% | 99.8285% | 99.8295% | 99.8285% | 99.8328% | 99.8312% | 99.8321% | 99.8319% | 99.8320% | 99.8319% |
| GATK | SNP | 9229 | 9169 | 9490 | 9364 | 9620 | 9403 | 9233 | 9367 | 9264 | 9080 |
| | InDel | 7669 | 6787 | 8338 | 7123 | 8520 | 9170 | 8606 | 8399 | 9500 | 7914 |
| Strelka 2 | SNP | 3419 | 3840 | 2983 | 2725 | 3726 | 3374 | 4033 | 3841 | 3713 | 4610 |
| | InDel | 5880 | 5461 | 5656 | 4690 | 9526 | 6367 | 6795 | 6436 | 10063 | 6133 |
| GATK and Strelka2 overlapping SNP+InDel | | NA | NA | NA | NA | 9083 | 7547 | 8046 | 7450 | 9727 | 7938 |
| Filter *rdr6* background | | NA | NA | NA | NA | 504 | 304 | 239 | 197 | 666 | 149 |
| Filter depth < 30 | | NA | NA | NA | NA | 491 | 293 | 226 | 187 | 649 | 135 |
| Filter Reference/Alternate allele reads ratio > 3 | | NA | NA | NA | NA | 203 | 162 | 124 | 94 | 293 | 66 |
| Overlap with Cas-OFFinder predicted off-targets | | NA | NA | NA | NA | 0 | 0 | 0 | 0 | 0 | 0 |

**Table S3.** High confidence variants (excluding the *AtPDS3* gRNA10 site) in CasΦ transgene free T2 albino seedlings.

| Chromosome | Position | Reference sequence | Variant sequence | T2 transgene negative plant | | | | | |
|---|---|---|---|---|---|---|---|---|---|
| | | | | *rdr6 vCasphi U6PDS3gR10* | | | *rdr6 nCasphi U6PDS3gR10* | | |
| | | | | line1 | line2 | line3 | line1 | line2 | line3 |
| chr1 | 6785692 | T | TC | ND* | Y* | ND | ND | ND | ND |
| chr1 | 7071325 | G | A | ND | ND | Y | ND | ND | ND |
| chr1 | 7636794 | G | A | ND | Y | ND | ND | ND | ND |
| chr1 | 9937635 | GTTGTA | G | ND | ND | ND | ND | Y | ND |
| chr1 | 11229308 | GTCTTTGTGTGAGC | G | ND | Y | ND | ND | ND | ND |
| chr1 | 11510229 | C | T | ND | ND | ND | ND | ND | Y |
| chr1 | 12571948 | A | T | ND | ND | Y | ND | ND | ND |
| chr1 | 13829012 | A | T | ND | ND | ND | ND | ND | Y |
| chr1 | 19941306 | TGCAATGAGGTTTTG | T | Y | ND | ND | ND | ND | ND |
| chr1 | 22302430 | GAAAGAAAC | G | ND | ND | ND | ND | ND | Y |
| chr1 | 26337186 | ATC | A | ND | ND | ND | Y | ND | ND |
| chr1 | 27602641 | A | T | Y | ND | ND | ND | ND | ND |
| chr1 | 28528239 | C | T | ND | ND | ND | Y | ND | ND |
| chr1 | 29180720 | CT | C | ND | ND | ND | ND | ND | Y |
| chr2 | 1123998 | G | A | ND | Y | ND | ND | ND | ND |
| chr2 | 1775712 | G | A | Y | ND | ND | ND | ND | ND |
| chr2 | 2396463 | A | G | ND | ND | ND | ND | ND | Y |
| chr2 | 3037107 | T | A | ND | Y | ND | ND | ND | ND |
| chr2 | 3189614 | T | A | Y | ND | ND | ND | ND | ND |
| chr2 | 3778595 | GA | G | ND | Y | ND | ND | ND | ND |
| chr2 | 4070245 | G | T | ND | ND | ND | ND | Y | ND |
| chr2 | 4473538 | G | A | ND | ND | ND | ND | Y | ND |
| chr2 | 4597486 | T | C | ND | ND | ND | Y | ND | ND |
| chr2 | 5939955 | A | T | ND | ND | ND | ND | ND | Y |
| chr2 | 9981681 | TTGATCAAGTAAATG ACATA | T | ND | ND | ND | ND | Y | ND |
| chr2 | 9982805 | TA | T | ND | ND | Y | ND | ND | ND |
| chr2 | 12499196 | T | TTA | ND | Y | ND | ND | ND | ND |
| chr2 | 13166249 | CAT | C | ND | Y | ND | ND | ND | ND |
| chr2 | 15371283 | TGAAG | T | Y | ND | ND | ND | ND | ND |
| chr2 | 15371288 | TCTAAATA | T | Y | ND | ND | ND | ND | ND |
| chr2 | 15371297 | TAA | T | Y | ND | ND | ND | ND | ND |
| chr2 | 15371300 | C | T | Y | ND | ND | ND | ND | ND |
| chr2 | 15371303 | A | T | Y | ND | ND | ND | ND | ND |
| chr2 | 15371304 | C | G | Y | ND | ND | ND | ND | ND |
| chr2 | 19035384 | T | C | ND | ND | ND | ND | ND | Y |
| chr2 | 19623291 | C | T | ND | ND | Y | ND | ND | ND |
| chr3 | 2788054 | A | T | Y | ND | ND | ND | ND | ND |
| chr3 | 4305186 | G | T | ND | ND | Y | ND | ND | ND |
| chr3 | 5085451 | TAGGGTCTA | T | ND | ND | ND | ND | Y | ND |

| | | | | | | | | | |
|---|---|---|---|---|---|---|---|---|---|
| chr3 | 8206493 | G | A | ND | Y | ND | ND | ND | ND |
| chr3 | 8609529 | C | T | ND | ND | ND | ND | Y | ND |
| chr3 | 12512717 | C | T | ND | ND | ND | Y | ND | ND |
| chr3 | 12613340 | A | T | ND | Y | ND | ND | ND | ND |
| chr3 | 14064862 | C | T | ND | ND | ND | Y | ND | ND |
| chr3 | 15021799 | A | T | ND | ND | ND | ND | Y | ND |
| chr3 | 16056216 | C | G | ND | ND | ND | ND | Y | ND |
| chr3 | 16378772 | TATACCTATACGA | T | ND | ND | ND | Y | ND | ND |
| chr3 | 16717520 | TGACGAGCTTGAG | T | ND | Y | ND | ND | ND | ND |
| chr3 | 17108050 | G | A | ND | ND | ND | Y | ND | ND |
| chr4 | 1843939 | G | A | ND | ND | ND | ND | ND | Y |
| chr4 | 2261579 | T | C | ND | ND | ND | ND | Y | ND |
| chr4 | 3360240 | C | T | Y | ND | ND | ND | ND | ND |
| chr4 | 5305862 | G | A | Y | ND | ND | ND | ND | ND |
| chr4 | 6581636 | C | CA | Y | ND | ND | ND | ND | ND |
| chr4 | 8633317 | TG | T | ND | ND | ND | ND | ND | Y |
| chr4 | 8801901 | G | A | ND | ND | ND | ND | ND | Y |
| chr4 | 10685528 | C | CA | ND | Y | ND | ND | ND | ND |
| chr4 | 15242376 | T | C | ND | Y | ND | ND | ND | ND |
| chr5 | 1309571 | A | G | ND | ND | ND | ND | ND | Y |
| chr5 | 2561456 | ATATGGTTTTGTTAACCGTG | A | ND | ND | ND | ND | Y | ND |
| chr5 | 6762332 | AAGTTTG | A | ND | ND | ND | ND | ND | Y |
| chr5 | 8524857 | T | A | ND | ND | ND | ND | Y | ND |
| chr5 | 9697518 | CCGTCAAAAACTATA | C | ND | Y | ND | ND | ND | ND |
| chr5 | 10036901 | G | A | ND | ND | ND | Y | ND | ND |
| chr5 | 10444172 | C | T | ND | Y | ND | ND | ND | ND |
| chr5 | 11347449 | A | T | ND | Y | Y | ND | ND | ND |
| chr5 | 12750772 | G | A | ND | ND | ND | Y | ND | ND |
| chr5 | 13934573 | GA | G | ND | ND | ND | ND | Y | ND |
| chr5 | 21062569 | C | T | ND | Y | ND | ND | ND | ND |
| chr5 | 23985255 | TA | T | ND | ND | ND | ND | ND | Y |
| chr5 | 26879986 | TTTAC | T | ND | Y | ND | ND | ND | ND |

* ND, not detected; Y, detected.

**Table S4.** Sequence of guide RNAs used.

| Purpose | CasΦ Guide RNA repeat sequence (common to all guides and on 5' of spacer sequence) | | |
|---|---|---|---|
| For plasmid vectors | GTCGGAACGCTCAACGATTGCCCCTCACGAGGGGAC | | |
| For RNPs | CAACGATTGCCCCTCACGAGGGGAC | | |
| **Guide RNA name** | **Guide RNA spacer sequence (Denoted in DNA sequence)** | **PAM** | **Direction relative to target gene** |
| *AtPDS3* gR8 | TTGTTCCGCAAAATAGCCCA | TCG | reverse |
| *AtPDS3* gR10 (20bp) | CAGTTGACAATCCAGCCAAT | TTG | reverse |
| *AtPDS3* gR10 (30bp) | CAGTTGACAATCCAGCCAATCCTGCAATTA | TTG | reverse |
| scramble control | GCGACACGACUCAUUAUA | none | not applicable |
| *FWA* gR1 | TCCCATTCAACATTCATACG | TTA | forward |
| *FWA* gR2 | TCGAAGCCCATACATCTTTC | TTA | forward |
| *FWA* gR3 | TGGGCCGAAGCCCATACATC | TTA | forward |
| *FWA* gR4 | TGGTTCTATACTAATATCAA | TTA | forward |
| *FWA* gR5 | ATATTAGTATAGAACCATAA | TTG | reverse |
| *FWA* gR6 | GTATAGAACCATAACAAAAG | TTA | reverse |
| *FWA* gR7 | CTAAATTTAGTAAAGAATCA | TTA | forward |
| *FWA* gR8 | GTAATCAATGGTTATTGTGA | TTA | reverse |
| *FWA* gR9 | TGAAATGAAATTTAACTTTT | TTG | reverse |
| *FWA* gR10 | GTTATCTAAATAAAACTAGG | TTA | forward |
| *Lc* gR1 | TGGACAGAGCTCCAAGTGAC | TTA | reverse |
| *Lc* gR2 | CTCGGTCACTTGGAGCTCTG | TTG | forward |
| *Lc* gR3 | GAGCTCTGTCCATAAATTAA | TTG | forward |
| *Lc* gR4 | TTGCCAACATAGAGTGTACG | TTA | forward |
| *Lc* gR5 | CCAACATAGAGTGTACGTGG | TTG | forward |
| *Lc* gR6 | CAGAAGCTAAACTCAACCAG | TTA | forward |
| *Lc* gR7 | GCTTCTGTAACACTACTGCT | TTA | reverse |
| *Lc* gR8 | TCTTTGGTGGAGCTCTGGTT | TTG | reverse |
| *Lc* gR9 | CTTGCAAATTGCATGCACGA | TTA | forward |
| *Lc* gR10 | CAAATTGCATGCACGAGCTA | TTG | forward |
| *Lc* gR11 | CATGCACGAGCTAGAATTAT | TTG | forward |

**Table S5.** Plasmids generated in this study.

| Plasmid name | Detailed information |
| --- | --- |
| HBT_pcoCASphi_version1 | cloning vector for sequence of FLAG-SV40NLS-CASphi-withIV2intron-nucleoplasminNLS (Version 1 arrangment) |
| HBT_pcoCASphi_version2 | cloning vector for sequence of CASphi-withIV2intron-2xSV40NLS-2xFLAG (Version 2 arrangment) |
| pC1300_pUB10_pcoCASphi_E9t_MCS_version1 | Binary vector with Arabidopsis codon optimized Casphi driven by UBQ10 promoter and RbcsE9 terminator. Design of NLS and Flag tag is indicated in Figure 1a version 1 plasmid. |
| pC1300_pUB10_pcoCASphi_E9t_MCS_version2 | Binary vector with Arabidopsis codon optimized Casphi driven by UBQ10 promoter and RbcsE9 terminator. Design of NLS and Flag tag is indicated in Figure 1a version 2 plasmid. |
| pC1300_pUB10_pcoCASphi_E9t_version1_U6_AtPDS3_gRNA10 | AtPDS3 guide RNA10 driven by AtU6-26 promoter was cloned into pC1300_pUB10_pcoCASphi_E9t_MCS_version1 plasmid |
| pC1300_pUB10_pcoCASphi_E9t_version2_U6_AtPDS3_gRNA10 | AtPDS3 guide RNA10 driven by AtU6-26 promoter was cloned into pC1300_pUB10_pcoCASphi_E9t_MCS_version2 plasmid |
| pC1300_pUB10_pcoCASphi_E9t_V2_CmYLCVp_35sT_A_form_AtPDS3_gRNA10 | single AtPDS3 gRNA10 driven by CmYLCV promoter and 35S terminator |
| pC1300_pUB10_pcoCASphi_E9t_V2_CmYLCVp_35sT_B_form_AtPDS3_gRNA10 | single AtPDS3 gRNA10 with an extra Casphi repeat sequence at the 3' end driven by CmYLCV promoter and 35S terminator |
| pC1300_pUB10_pcoCASphi_E9t_V2_CmYLCVp_35sT_C_form_AtPDS3_gRNA10 | triple AtPDS3 gRNA10 array driven by CmYLCV promoter and 35S termnator |
| pC1300_pUB10_pcoCASphi_E9t_V2_2x35Sp_HSP18t_A_form_AtPDS3_gRNA10 | single AtPDS3 gRNA10 driven by 2x35S promoter and HSP18 terminator |
| pC1300_pUB10_pcoCASphi_E9t_V2_2x35Sp_HSP18t_B_form_AtPDS3_gRNA10 | single AtPDS3 gRNA10 with an extra Casphi repeat sequence at the 3' end driven by 2x35S promoter and HSP18 terminator |
| pC1300_pUB10_pcoCASphi_E9t_V2_2x35Sp_HSP18t_C_form_AtPDS3_gRNA10 | triple AtPDS3 gRNA10 driven by 2x35S promoter and HSP18 terminator |
| pC1300_pUB10_pcoCASphi_E9t_V2_pUB10_E9t_A_form_AtPDS3_gRNA10 | single AtPDS3 gRNA10 driven by UBQ10 promoter and Rbcs E9 terminator |
| pC1300_pUB10_pcoCASphi_E9t_V2_pUB10_E9t_B_form_AtPDS3_gRNA10 | single AtPDS3 gRNA10 with an extra Casphi repeat sequence at the 3' end driven by UBQ10 promoter and Rbcs E9 terminator |
| pC1300_pUB10_pcoCASphi_E9t_V2_pUB10_E9t_C_form_AtPDS3_gRNA10 | triple AtPDS3 gRNA10 driven by UBQ10 promoter and Rbcs E9 terminator |
| pC1300_pUB10_pcoCASphi_E9t_V2_CmYLCVp_35sT_A_form_AtPDS3_gRNA10_30bp_spacer | single AtPDS3 gRNA10 with 30bp spacer sequence driven by CmYLCV promoter and 35S terminator |
| pC1300_pUB10_pcoCASphi_E9t_V2_CmYLCVp_35sT_C_form_AtPDS3_gRNA10_30bp_spacer | triple AtPDS3 gRNA10 with 30bp spacer sequence array driven by CmYLCV promoter and 35S termnator |
| pC1300_pUB10_pcoCASphi_E9t_V2_2x35Sp_HSP18t_C_form_AtPDS3_gRNA10_30bp_spacer | triple AtPDS3 gRNA10 with 30bp spacer sequence driven by 2x35S promoter and HSP18 terminator |
| pC1300_pUB10_pcoCASphi_E9t_V2_pUB10_E9t_C_form_AtPDS3_gRNA10_30bp_spacer | triple AtPDS3 gRNA10 with 30bp spacer sequence driven by UBQ10 promoter and Rbcs E9 terminator |
| pC1300_pUB10_pcoCASphi_E9t_V2_CmYLCVp_35sT_ribozyme_AtPDS3_gRNA10 | single AtPDS3 gRNA10 flanked by ribozymes driven by CmYLCV promoter and 35S terminator |
| pC1300_pUB10_pcoCASphi_E9t_V2_2x35Sp_HSP18t_ribozyme_AtPDS3_gRNA10 | single AtPDS3 gRNA10 flanked by ribozymes driven by 2x35S promoter and HSP18 terminator |
| pC1300_pUB10_pcoCASphi_E9t_V2_pUB10_E9t_ribozyme_AtPDS3_gRNA10 | single AtPDS3 gRNA10 flanked by ribozymes driven by UBQ10 promoter and Rbcs E9 terminator |

| | |
|---|---|
| pC1300_pUB10_pco-vCASphi_E9t_MCS_version2 | Binary vector with Arabidopsis codon optimized vCasphi driven by UBQ10 promoter and RbcsE9 terminator. Design of NLS and Flag tag is same to the pC1300_pUB10_pcoCASphi_E9t_MCS_version2 plasmid. |
| pC1300_pUB10_pco-nCASphi_E9t_MCS_version2 | Binary vector with Arabidopsis codon optimized nCasphi driven by UBQ10 promoter and RbcsE9 terminator. Design of NLS and Flag tag is same to the pC1300_pUB10_pcoCASphi_E9t_MCS_version2 plasmid. |
| pC1300_pUB10_pcoCASphi_E9t_V2_U6_AtPDS3_gRNA8 | AtPDS3 guide RNA8 driven by AtU6-26 promoter was cloned into pC1300_pUB10_pcoCASphi_E9t_MCS_version2 plasmid |
| pC1300_pUB10_pcoCASphi_E9t_V2_U6_FWA_gRNA1 | FWA guide RNA1 driven by AtU6-26 promoter was cloned into pC1300_pUB10_pcoCASphi_E9t_MCS_version2 plasmid |
| pC1300_pUB10_pcoCASphi_E9t_V2_U6_FWA_gRNA4 | FWA guide RNA4 driven by AtU6-26 promoter was cloned into pC1300_pUB10_pcoCASphi_E9t_MCS_version2 plasmid |
| pC1300_pUB10_pcoCASphi_E9t_V2_U6_FWA_gRNA5 | FWA guide RNA5 driven by AtU6-26 promoter was cloned into pC1300_pUB10_pcoCASphi_E9t_MCS_version2 plasmid |
| pC1300_pUB10_pcoCASphi_E9t_V2_U6_FWA_gRNA6 | FWA guide RNA6 driven by AtU6-26 promoter was cloned into pC1300_pUB10_pcoCASphi_E9t_MCS_version2 plasmid |
| pC1300_pUB10_pco-vCASphi_E9t_V2_U6_AtPDS3_gRNA8 | AtPDS3 guide RNA8 driven by AtU6-26 promoter was cloned into pC1300_pUB10_pco-vCASphi_E9t_MCS_version2 plasmid |
| pC1300_pUB10_pco-vCASphi_E9t_V2_U6_AtPDS3_gRNA10 | AtPDS3 guide RNA10 driven by AtU6-26 promoter was cloned into pC1300_pUB10_pco-vCASphi_E9t_MCS_version2 plasmid |
| pC1300_pUB10_pco-vCASphi_E9t_V2_U6_FWA_gRNA1 | FWA guide RNA1 driven by AtU6-26 promoter was cloned into pC1300_pUB10_pco-vCASphi_E9t_MCS_version2 plasmid |
| pC1300_pUB10_pco-vCASphi_E9t_V2_U6_FWA_gRNA4 | FWA guide RNA4 driven by AtU6-26 promoter was cloned into pC1300_pUB10_pco-vCASphi_E9t_MCS_version2 plasmid |
| pC1300_pUB10_pco-vCASphi_E9t_V2_U6_FWA_gRNA5 | FWA guide RNA5 driven by AtU6-26 promoter was cloned into pC1300_pUB10_pco-vCASphi_E9t_MCS_version2 plasmid |
| pC1300_pUB10_pco-vCASphi_E9t_V2_U6_FWA_gRNA6 | FWA guide RNA6 driven by AtU6-26 promoter was cloned into pC1300_pUB10_pco-vCASphi_E9t_MCS_version2 plasmid |
| pC1300_pUB10_pco-nCASphi_E9t_V2_U6_AtPDS3_gRNA8 | AtPDS3 guide RNA8 driven by AtU6-26 promoter was cloned into pC1300_pUB10_pco-nCASphi_E9t_MCS_version2 plasmid |
| pC1300_pUB10_pco-nCASphi_E9t_V2_U6_AtPDS3_gRNA10 | AtPDS3 guide RNA10 driven by AtU6-26 promoter was cloned into pC1300_pUB10_pco-nCASphi_E9t_MCS_version2 plasmid |
| pC1300_pUB10_pco-nCASphi_E9t_V2_U6_FWA_gRNA1 | FWA guide RNA1 driven by AtU6-26 promoter was cloned into pC1300_pUB10_pco-nCASphi_E9t_MCS_version2 plasmid |
| pC1300_pUB10_pco-nCASphi_E9t_V2_U6_FWA_gRNA4 | FWA guide RNA4 driven by AtU6-26 promoter was cloned into pC1300_pUB10_pco-nCASphi_E9t_MCS_version2 plasmid |
| pC1300_pUB10_pco-nCASphi_E9t_V2_U6_FWA_gRNA5 | FWA guide RNA5 driven by AtU6-26 promoter was cloned into pC1300_pUB10_pco-nCASphi_E9t_MCS_version2 plasmid |
| pC1300_pUB10_pco-nCASphi_E9t_V2_U6_FWA_gRNA6 | FWA guide RNA6 driven by AtU6-26 promoter was cloned into pC1300_pUB10_pco-nCASphi_E9t_MCS_version2 plasmid |

| | |
|---|---|
| pC1300_pUB10_pco-vCASphi_E9t_V2_CmYLCVp_35sT_ribozyme_AtPDS3_gRNA10 | the cassette of single AtPDS3 guide RNA10 flanded by ribozymes driven by the CmYLCV promoter and 35S terminator was cloned into the pC1300_pUB10_pco-vCASphi_E9t_MCS_version2 vector |
| pC1300_pUB10_pco-vCASphi_E9t_V2_pUB10_E9t_ribozyme_AtPDS3_gRNA10 | the cassette of single AtPDS3 guide RNA10 flanded by ribozymes driven by the UBQ10 promoter and Rbcs E9 terminator was cloned into the pC1300_pUB10_pco-vCASphi_E9t_MCS_version2 vector |
| pC1300_pUB10_pco-nCASphi_E9t_V2_CmYLCVp_35sT_ribozyme_AtPDS3_gRNA10 | the cassette of single AtPDS3 guide RNA10 flanded by ribozymes driven by the CmYLCV promoter and 35S terminator was cloned into the pC1300_pUB10_pco-nCASphi_E9t_MCS_version2 vector |
| pC1300_pUB10_pco-nCASphi_E9t_V2_pUB10_E9t_ribozyme_AtPDS3_gRNA10 | the cassette of single AtPDS3 guide RNA10 flanded by ribozymes driven by the UBQ10 promoter and Rbcs E9 terminator was cloned into the pC1300_pUB10_pco-nCASphi_E9t_MCS_version2 vector |

**Table S6.** Sequence of primers and synthesized double stranded DNA.

| For amplicon sequencing: | | |
|---|---|---|
| **Oligo name** | **Oligo sequence** | **Purpose and details** |
| 20685 | ACACTCTTTCCCTACACGACGCTCTTCCGATCTCATGGCTGGCAAAAGTCCAATAGCA | AtPDS3 gR8 amplicon step1 FW |
| 20686 | GTGACTGGAGTTCAGACGTGTGCTCTTCCGATCTACTGGTCAAGGCAAGACGATATAACT | AtPDS3 gR8 amplicon step1 RV |
| 20681 | ACACTCTTTCCCTACACGACGCTCTTCCGATCTGTACCTTTCCACCAAGAACATCTCT | AtPDS3 gR10 amplicon step1 FW |
| 20682 | GTGACTGGAGTTCAGACGTGTGCTCTTCCGATCTTCCTCGTCCTGCTAAGCCTTTGA | AtPDS3 gR10 amplicon step1 RV |
| 21405 | ACACTCTTTCCCTACACGACGCTCTTCCGATCTTTCGTTCTTGTGTCATGTAATAGATTACT | FWA gR10 amplicon step1 FW |
| 21406 | GTGACTGGAGTTCAGACGTGTGCTCTTCCGATCTGCCCATAACTCTTTGATATTAGTATAGA | FWA gR10 amplicon step1 RV |
| 21407 | ACACTCTTTCCCTACACGACGCTCTTCCGATCTAGGCCATCCATGGATGGTTTCA | FWA gR7, gR8, gR9 amplicon step1 FW |
| 21408 | GTGACTGGAGTTCAGACGTGTGCTCTTCCGATCTGAATATATGAGATTCTCGACGGAAAGA | FWA gR7, gR8, gR9 amplicon step1 RV |
| 21409 | ACACTCTTTCCCTACACGACGCTCTTCCGATCTGAATATATGAGATTCTCGACGGAAAGA | FWA gR4, gR5, gR6 amplicon step1 FW |
| 21410 | GTGACTGGAGTTCAGACGTGTGCTCTTCCGATCTAGGCCATCCATGGATGGTTTCA | FWA gR4, gR5, gR6 amplicon step1 RV |
| 21411 | ACACTCTTTCCCTACACGACGCTCTTCCGATCTCGGAAAGATGTATGGGCTTCGAT | FWA gR3 amplicom step1 C12J FW, pair with primer 21410 for PCR reaction |
| 21412 | ACACTCTTTCCCTACACGACGCTCTTCCGATCTATAGCACTTGGACCAATGGCGAA | FWA gR1 amplicon step1 FW, pair with primer 21414 for PCR reaction |
| 21413 | ACACTCTTTCCCTACACGACGCTCTTCCGATCTCGTATGAATGTTGAATGGGATAAGGT | FWA gR2 amplicon step1 FW, pair with primer 21414 for PCR reaction |
| 21414 | GTGACTGGAGTTCAGACGTGTGCTCTTCCGATCTAGAATCAATTGGGTTTAGTGTTTACTTGT | FWA gR1, gR2 amplicon step1 RV |
| 23997 | ACACTCTTTCCCTACACGACGCTCTTCCGATCTGACGGGGTAGATGCCTACAGA | casphi lc gR1,gR2,gR3 amplicon step1 FW |
| 23998 | GTGACTGGAGTTCAGACGTGTGCTCTTCCGATCTTATAGCGTCAGAGAACTTAGATCTGA | casphi lc gR1,gR2,gR3 ,gR6,gR7,gR8 amplicon step1 RV |
| 23999 | ACACTCTTTCCCTACACGACGCTCTTCCGATCTAGTGACCGAGCAAGACGGTGA | casphi lc gR6,gR7,gR8 amplicon step1 FW |
| 24000 | ACACTCTTTCCCTACACGACGCTCTTCCGATCTGCTCCTCACTAGCTACCAAGAG | casphi lc gR4,gR5 amplicon step1 FW |
| ZL21 | GTGACTGGAGTTCAGACGTGTGCTCTTCCGATCTCTACTGATCATCAGACGATGCCT | casphi lc gR4,gR5 amplicon step1 RV |
| ZL22 | ACACTCTTTCCCTACACGACGCTCTTCCGATCTCACACAGTATCAGCTGGCACA | casphi lc gR9,gR10,gR11 amplicon step1 FW |
| ZL23 | GTGACTGGAGTTCAGACGTGTGCTCTTCCGATCTCTGGTTGAGTGTCTGAAATGGAC | casphi lc gR9,gR10,gR11 amplicon step1 RV |

| | | |
|---|---|---|
| **For cloning and genotyping:** | | |
| **Oligo name** | **Oligo sequence** | **Purpose and details** |
| 20478 | AGCAGCTGGAACTCCGTGGA | FW primer to amplify HBT backbone for HBT_pcoCASphi version1 |
| 20479 | AAGAGACCAGCTGCTACCAAGA | RV primer to amplify HBT backbone for HBT_pcoCASphi version1 |
| 20497 | GGATCCACGGAGCAAGGGGA | FW primer to amplify HBT backbone for HBT_pcoCASphi version2 |
| 20498 | TGACTGCAGATCGTTCAAACATTTGGCA | RV primer to amplify HBT backbone for HBT_pcoCASphi version2 |
| 20501 | TCCACGGAGTTCCAGCTGCTATGCCGAAGCCCGCCGTCGA | FW primer to amplify pcoCASphi 1st fragment for in-fusion reaction to generate HBT_pcoCASphi_version1 |
| 20507 | CTTGCTCCGTGGATCCATGCCGA | FW primer to amplify pcoCASphi 1st fragment for in-fusion reaction to generate HBT_pcoCASphi_version2 |
| 20502 | CTGGCGTTTCTTAGTAATCCTCTTACGT | RV primer to amplify pcoCASphi 1st fragment for in-fusion reaction to generate HBT_pcoCASphi_version1 and HBT_pcoCASphi_version2 |
| 20503 | GGATTACTAAGAAACGCCAGGTAAGTTTCTGCTTCTACCTTTGA | FW primer to amplify IV2 intron for in-fusion reaction to generate HBT_pcoCASphi_version1 and HBT_pcoCASphi_version2 |
| 20504 | TGGGAGCAATAGTCCTCCACCTGCACATCAACAAATTTTGGTCA | RV primer to amplify IV2 intron for in-fusion reaction to generate HBT_pcoCASphi_version1 and HBT_pcoCASphi_version2 |
| 20505 | GTGGAGGACTATTGCTCCCAAGGA | FW primer to amplify pcoCASphi 2nd fragment for in-fusion reaction to generate HBT_pcoCASphi_version1 and HBT_pcoCASphi_version2 |
| 20506 | TTGGTAGCAGCTGGTCTCTTGGACGTTTGGGACGGCTCTTGA | RV primer to amplify pcoCASphi 2nd fragment for in-fusion reaction to generate HBT_pcoCASphi_version1 |
| 20508 | GCCAAATGTTTGAACGATCTGCAGTCACT | RV primer to amplify pcoCASphi 2nd fragment for in-fusion reaction to generate HBT_pcoCASphi_version2 |

| | | |
|---|---|---|
| 20544 | ATCACTAGTATCCTAGGAAGGTACCAGTCTAGCTCAACAGAGCTTTTAACCCA | FW primer to amplify pUBQ10 for in-fusion reaction to generate pC1300_pUB10_pcoCASphi_E9t_MCS_version1 and pC1300_pUB10_pcoCASphi_E9t_MCS_version2 plasmids |
| 20545 | TCATCATCCTTGTAATCCATCTGTTAATCAGAAAAACTCAGATTAATCGA | RV primer to amplify pUBQ10 for in-fusion reaction to generate pC1300_pUB10_pcoCASphi_E9t_MCS_version1 |
| 20552 | TCGACGGCGGGCTTCGGCATCTGTTAATCAGAAAAACTCAGATTAATCGA | RV primer to amplify pUBQ10 for in-fusion reaction to generate pC1300_pUB10_pcoCASphi_E9t_MCS_version2 |
| 20548 | TGAGTTTTTCTGATTAACAGATGGATTACAAGGATGATGATGATAAGGA | FW primer to amplify 2xFLAG-SV40-Casphi-2-IV2intron-nucleoplasminNLS for in-fusion reaction to generate pC1300_pUB10_pcoCASphi_E9t_MCS_version1 |
| 20550 | TGAGTTTTTCTGATTAACAGATGCCGAAGCCCGCCGTCGAATCA | FW primer to amplify Casphi-2-IV2intron-2xSV40-2xFLAG for in-fusion reaction to generate pC1300_pUB10_pcoCASphi_E9t_MCS_version2 |
| 20549 | ATGATACGAACGAAAGCTCTTCACTTCTTCTTCTTAGCCTGTCCA | RV primer to amplify 2xFLAG-SV40-Casphi-2-IV2intron-nucleoplasminNLS for in-fusion reaction to generate pC1300_pUB10_pcoCASphi_E9t_MCS_version1 |
| 20551 | ATGATACGAACGAAAGCTCTTCACTTGTCGTCGTCATCCTTATAGT | RV primer to amplify Casphi-2-IV2intron-2xSV40-2xFLAG for in-fusion reaction to generate pC1300_pUB10_pcoCASphi_E9t_MCS_version2 |
| 20546 | AGGCTAAGAAGAAGAAGTGAAGAGCTTTCGTTCGTATCATCGGT | FW primer to amplify RbcS E9 terminator for in-fusion reaction to generate pC1300_pUB10_pcoCASphi_E9t_MCS_version1 |
| 20553 | AGGATGACGACGACAAGTGAAGAGCTTTCGTTCGTATCATCGGTTTCGA | FW primer to amplify RbcS E9 terminator for in-fusion reaction to generate pC1300_pUB10_pcoCASphi_E9t_MCS_version2 |

| | | |
|---|---|---|
| 20547 | CAGCTATGACCATGATTACGAATTCGTTGTCAATCA ATTGGCAAGTCATAAAATGCA | RV primer to amplify RbcS E9 terminator for in-fusion reaction to generate pC1300_pUB10_pcoCASphi_E9t_MCS_version1 and pC1300_pUB10_pcoCASphi_E9t_MCS_version2 plasmids. |
| 14444 | TTCCTAGGATACTAGAAGCTTCGTTGAACAACGGA | FW primer to amplify AtU6-26 gRNA cassette for in-fusion reaction |
| 20665 | TGCAGGTCGACTCTAGATCACTAGTGATCAGATGCA GAGAGACT | RV primer to amplify AtU6-26 gRNA cassette for in-fusion reaction |
| 20730 | TTGTTCCGCAAAATAGCCCATTTTTTTTGCCATTCTTT TCAAGCTCCA | AtPDS3 gRNA8 FW |
| 20731 | TGGGCTATTTTGCGGAACAAGTCCCCTCGTGAGGG GCAATCGTTGA | AtPDS3 gRNA8 RV |
| 20734 | CAGTTGACAATCCAGCCAATTTTTTTTTGCCATTCTTT TCAAGCTCCA | AtPDS3 gRNA10 FW |
| 20735 | ATTGGCTGGATTGTCAACTGGTCCCCTCGTGAGGGG CAATCGTTGA | AtPDS3 gRNA10 RV |
| 21950 | TCCCATTCAACATTCATACGTTTTTTTTGCCATTCTTT TCAAGCTCCATTGTCA | FWA gRNA1 FW |
| 21949 | CGTATGAATGTTGAATGGGAGTCCCCTCGTGAGGG GCAATCGTTGAGCGTTCCGA | FWA gRNA1 RV |
| 21952 | TGGTTCTATACTAATATCAATTTTTTTTGCCATTCTTT TCAAGCTCCATTGTCA | FWA gRNA4 FW |
| 21951 | TTGATATTAGTATAGAACCAGTCCCCTCGTGAGGGG CAATCGTTGAGCGTTCCGA | FWA gRNA4 RV |
| 21954 | ATATTAGTATAGAACCATAATTTTTTTTGCCATTCTTT TCAAGCTCCATTGTCA | FWA gRNA5 FW |
| 21953 | TTATGGTTCTATACTAATATGTCCCCTCGTGAGGGG CAATCGTTGAGCGTTCCGA | FWA gRNA5 RV |
| 21956 | GTATAGAACCATAACAAAAGTTTTTTTTGCCATTCTT TTCAAGCTCCATTGTCA | FWA gRNA6 FW |
| 21955 | CTTTTGTTATGGTTCTATACGTCCCCTCGTGAGGGGC AATCGTTGAGCGTTCCGA | FWA gRNA6 RV |
| 21276 | GACTGGTACCTTCCTAGGATACTAGTGGCAGACATA CTGTCCCACA | CmYLCV promoter FW |
| 21277 | GACTTTGATAGCTTGCTGAGGCAAGCTTAGCTCTTA CCTGTTTTCGT | CmYLCV promoter RV |
| 21278 | TTCGATAATTCCTTAATTAACTCGAGTTTCTCCATAA TAATGTGTGA | 35S terminator FW |
| 21386 | CTGCAGGTCGACTCTAGATCACTAGTTAATTCGGGG GATCTGGATTTTAGTACT | 35S terminator RV |
| 21440 | CTGTTGAGCTAGACTGGTACCTTCCTAGGATACTAG GCCAGTGCCAAGCTTG | 2x35S promoter FW |

| | | |
|---|---|---|
| 21388 | GACTTTGATAGCTTGCTGAGGCGGGATCCTCTAGAG TCGAGGT | 2x35S promoter RV |
| 21389 | TTCGATAATTCCTTAATTAAATATGAAGATGAAGAT GAAATATTTGGTGTGT | HSP18.2 terminator FW |
| 21390 | CTGCAGGTCGACTCTAGATCACTAGTCTTATCTTTAA TCATATTCCATAGTCCATACCA | HSP18.2 terminator RV |
| 21451 | TTGAGCTAGACTGGTACCTTCCTAGGATACTAGAAG TTGTAATGAGTTGCTGGCCTCTCT | TBSinsulator-UBQ10 promoter FW |
| 21392 | GACTTTGATAGCTTGCTGAGGCCTGTTAATCAGAAA AACTCAGATTAATCGACA | UBQ10 promoter RV |
| 21393 | TTCGATAATTCCTTAATTAAAGAGCTTTCGTTCGTAT CATCGGT | Rbcs E9 terminator FW |
| 21394 | CTGCAGGTCGACTCTAGATCACTAGTGTTGTCAATC AATTGGCAAGTCATAAAATGCA | Rbcs E9 terminator RV |
| single PDS3 gRNA10 FW | GCCTCAGCAAGCTATCAAAGTCGGAACGCTCAACGA TTGCCCCTCACGAGGGGACCAGTT | single PDS3 gRNA10 FW to clone into PolII promoter gRNA cassette |
| single PDS3 gRNA10 RV | TTAATTAAGGAATTATCGAAATTGGCTGGATTGTCA ACTGGTCCCCTCGTGAGG | single PDS3 gRNA10 RV to clone into PolII promoter gRNA cassette |
| single PDS3 gR10 + repeat FW | GCCTCAGCAAGCTATCAAAGTCGGAACGCTCAACGA TTGCCCCTCACGAGGGGACCAGTTGACAATCCAGCC AAT | single PDS3 gR10 + repeat FW to clone into PolII promoter gRNA cassette |
| single PDS3 gR10 + repeat RV | TTAATTAAGGAATTATCGAAGTCCCCTCGTGAGGGG CAATCGTTGAGCGTTCCGACATTGGCTGGATTGTCA ACTG | single PDS3 gR10 + repeat RV to clone into PolII promoter gRNA cassette |
| triple PDS3 gR10 FW | GCCTCAGCAAGCTATCAAAGTCGGAACGCTCAACGA TTGCCCCTCACGAGGGGACCAGTTGACAATCCAGCC AATGTCGGAACGCTCAACGATTGCCCCTCACGAGG GGACCAGTTGACAATCCAGCCAATGTCGGAACGCTC AACGATTGCCCCTCACGAGGGGACCAGTTGACAATC CAGCCAAT | triple PDS3 gR10 FW to clone into PolII promoter gRNA cassette |
| triple PDS3 gR10 RV | TTAATTAAGGAATTATCGAAGTCCCCTCGTGAGGGG CAATCGTTGAGCGTTCCGACATTGGCTGGATTGTCA ACTGGTCCCCTCGTGAGGGGCAATCGTTGAGCGTTC CGACATTGGCTGGATTGTCAACTGGTCCCCTCGTGA GGGGCAATCGTTGAGCGTTCCGACATTGGCTGGATT GTCAACTG | triple PDS3 gR10 RV to clone into PolII promoter gRNA cassette |
| 21757 | AAGCTTGCCTCAGCAAGCTATCAAAGTCGGAACGCT CAACGATTG | PolII gRNA cassette fw (to clone the gRNA of 30bp spacer into the PolII gRNA cassette) |
| 21760 | AACTCGAGTTAATTAAGGAATTATCGAATAATTGCA GGATTGGCTGGATTGTCAACTGGT | PolII gRNA cassette AtPDS3 gRNA10 30bp RV |

| | | |
|---|---|---|
| 21763 | TGCCTCAGCAAGCTATCAAAGTCGGAACGCTCAACGATTGCCCCTCACGAGGGGACCAGTTGACAATCCAGCCAATCCTGCAATTAGTCGGAACGCTCAACGATTGCCCCTCACGAGGGGACCAGTTGACAATCCAGCCAATCCTGCAATTAGTCGGAACGCTCAACGATTGCCCCTCACGAGGGGACCAGTTGACAATC | 30bp spacer PDS3 triple gR10 array fw |
| 21764 | AGTTAATTAAGGAATTATCGAAGTCCCCTCGTGAGGGGCAATCGTTGAGCGTTCCGACTAATTGCAGGATTGGCTGGATTGTCAACTGGTCCCCTCGTGAGGGGCAATCGTTGAGCGTTCCGACTAATTGCAGGATTGGCTGGATTGTCAACTGGTCCCCTCGTGAGGGGCAATCGTTGAGCGTTCCGACTAATTGCAGG | 30bp spacer PDS3 triple gR10 array rv |
| 21732 | TAAGAGCTAAGCTTGCCTCAGCTCCGACCTGATGAGTCCGTGAGGACGAAACGAGTAAGCTCGTCGTCGGAACGCTCAACGATTGCCCCTCACGAGGGGACCAGTTGACAATCCAGCCAATG | PDS3 gR10 + ribozyme fw |
| 21733 | TGGAGAAACTCGAGTTAATTAAGTCCCATTCGCCATGCCGAAGCATGTTGCCCAGCCGGCGCCAGCGAGGAGGCTGGGACCATGCCGGCCATTGGCTGGATTGTCAACTGGT | PDS3 gRNA10 + ribozyme rv |
| 21847 | TCCTCCTCACCTGAAGATCCGGCCTTCTCATTTCGAAGCTGT | vCasphi mutation RV |
| 21848 | GGCCGGATCTTCAGGTGAGGAGGAGGTAGCTACAAATGA | vCasphi mutation FW |
| 21849 | TCTTACGTCTATGACTACCCAATCGGT | vCasphi and nCasphi Fragment2 RV |
| 21851 | AGCTGCAGCTCGAGCGGCGTTTATTGCAGCCAATCTAGCTCGGGCCTTCTCA | nCasphi mutation RV |
| 21852 | GCTGCAATAAACGCCGCTCGAGCTGCAGCTGGATTGCCGGAAATCAAGGCCGAGGA | nCasphi mutation FW |
| 20505 | GTGGAGGACTATTGCTCCCAAGGA | Casphi genotyping FW |
| 20639 | TCCAGAGCTCTGACCTCTGCT | Casphi genotyping RV |
| 21403 | AAGGAGTCATTTTTCACTAAGCATATAGA | FWA fragment amplification FW for in vitro RNP cleavage substrate |
| 21404 | CATTTCTAGTGTCTCGACAACGAACA | FWA fragment amplification RV for in vitro RNP cleavage substrate |
| **For real-time quantitative PCR:** | | |
| **Oligo name** | **Oligo sequence** | **Purpose and details** |
| 11859 | GTATGAGTTGCTTCTCCAGCAAAG | IPP2 QPCR FW |
| 11860 | GAGGATGGCTGCAACAAGTGT | IPP2 QPCR RV |
| 21056 | GGTCGGAACGCTCAACGATTG | CASphi gRNA QPCR FW |
| 21059 | ATTGGCTGGATTGTCAACTGGTC | CASphi AtPDS3gR10 QPCR RV |
| **DNA synthesized:** | | |

| **DNA name** | **sequence** |
|---|---|
| CASphi-2-2xSV40NLS-2xFLAG | CTTGCTCCGTGGATCCATGCCGAAGCCCGCCGTCGA<br>ATCAGAGTTTTCCAAAGTCCTCAAGAAACACTTTCCT<br>GGGGAGCGTTTTAGGTCTAGCTATATGAAGAGGGG<br>GGGTAAAATTCTGGCAGCACAAGGCGAGGAAGCTG<br>TAGTGGCGTACTTGCAGGGAAAGAGTGAGGAGGA<br>ACCGCCGAATTTTCAGCCGCCGGCGAAGTGCCACGT<br>GGTCACCAAAAGCAGGGATTTCGCAGAATGGCCCA<br>TAATGAAAGCCTCTGAAGCCATACAGAGGTACATCT<br>ATGCGCTCAGCACTACAGAGCGAGCTGCCTGCAAA<br>CCGGGTAAGAGCTCAGAAAGTCACGCGGCCTGGTT<br>CGCGGCTACAGGGGTGAGCAATCACGGCTATTCTC<br>ATGTACAAGGTCTTAACCTGATCTTTGACCACACGCT<br>AGGACGATACGATGGCGTTTTAAAGAAAGTACAGC<br>TTCGAAATGAGAAGGCCCGAGCTAGATTGGAAAGC<br>ATAAACGCCTCACGAGCTGATGAAGGATTGCCGGA<br>AATCAAGGCCGAGGAGGAGGAGGTAGCTACAAAT<br>GAAACAGGTCATCTACTACAGCCGCCAGGCATAAAC<br>CCATCATTCTACGTCTACCAGACCATATCTCCGCAGG<br>CTTACCGACCAAGGGACGAAATAGTGTTACCACCCG<br>AGTACGCCGGTTACGTCAGGGATCCGAACGCTCCG<br>ATTCCACTGGGCGTGGTCAGGAACCGTTGTGACATA<br>CAGAAGGGTTGCCCGGGATATATACCCGAGTGGCA<br>GAGGGAAGCTGGTACGGCAATTAGTCCCAAGACAG<br>GAAAAGCAGTGACGGTTCCAGGACTTAGCCCGAAG<br>AAGAATAAACGTATGCGTAGGTACTGGAGGTCAGA<br>AAAGGAGAAGGCTCAAGATGCACTTCTCGTAACTGT<br>AAGGATAGGTACCGATTGGGTAGTCATAGACGTAA<br>GAGGATTACTAAGAAACGCCAGGTGGAGGACTATT<br>GCTCCCAAGGACATAAGTTTAAATGCACTTCTAGAT<br>TTATTTACCGGTGATCCAGTCATCGATGTCAGACGA<br>AACATCGTGACCTTCACCTATACCTTGGACGCTTGC<br>GGAACTTATGCTAGAAAATGGACTCTCAAGGGAAA<br>ACAGACAAAAGCAACCTTAGATAAACTGACAGCGA<br>CACAAACTGTGGCCTTAGTTGCTATAGATCTGGGAC<br>AAACAAACCCAATTAGCGCGGGTATCAGTCGTGTCA<br>CACAGGAGAACGGGGCCCTCCAGTGCGAACCGCTT<br>GATCGTTTTACATTGCCTGATGACCTTTTGAAAGATA<br>TTTCTGCGTACCGAATTGCATGGGACCGTAACGAGG<br>AGGAACTCAGGGCCAGATCCGTTGAGGCACTCCCA<br>GAGGCACAACAAGCAGAGGTCAGAGCTCTGGACGG<br>GGTCTCCAAAGAGACCGCGCGTACACAGTTGTGCG<br>CGGACTTCGGTCTGGACCCAAAGCGACTACCGTGG<br>GATAAAATGAGTAGCAATACCACGTTCATAAGCGA<br>GGCGCTCCTTTCCAACAGCGTATCCCGTGACCAAGT<br>ATTCTTTACCCCGGCCCCAAAGAAAGGAGCCAAGAA<br>GAAAGCACCGGTGGAAGTGATGCGAAAAGACAGG |

| | |
|---|---|
| | ACATGGGCGCGAGCGTACAAACCACGACTCTCAGT<br>AGAAGCACAAAAGTTGAAAAATGAGGCTCTTTGGG<br>CTTTGAAGCGTACCTCTCCAGAATATCTAAAGTTGTC<br>ACGACGTAAAGAAGAATTGTGTAGGAGGTCCATTA<br>ATTACGTGATAGAAAAAACTAGAAGGCGAACCCAA<br>TGCCAAATTGTGATCCCCGTTATAGAAGATCTAAAT<br>GTCCGATTTTTCCACGGGTCTGGCAAACGACTCCCG<br>GGCTGGGATAACTTCTTCACGGCAAAGAAGGAAAA<br>CCGATGGTTTATCCAAGGGCTACATAAGGCGTTTTC<br>TGACTTAAGGACCCACCGATCCTTCTACGTGTTCGA<br>GGTGCGTCCAGAGAGAACATCAATCACCTGTCCGA<br>AATGCGGGCACTGTGAAGTGGGGAACAGGGATGG<br>AGAAGCCTTTCAGTGTCTCAGTTGCGGTAAGACATG<br>CAACGCAGATCTGGACGTAGCGACACATAACTTGAC<br>TCAAGTGGCGCTCACCGGCAAAACAATGCCGAAGA<br>GAGAGGAGCCTAGAGATGCACAAGGGACAGCCCCC<br>GCGAGAAAGACGAAGAAAGCTAGCAAGTCAAAGG<br>CACCCCCGGCTGAACGTGAGGATCAGACTCCCGCTC<br>AAGAGCCGTCCCAAACGTCCGGATCCGGACCGAAG<br>AAAAAGCGAAAGGTAGAGGATCCTAAAAAGAAGC<br>GTAAAGTCTCCTTGGGTTCTGGCTCCGACTATAAGG<br>ATGACGATGACAAAGACTATAAGGATGACGACGAC<br>AAGTGACTGCAGATCGTTCAAACATTTGGC |
